# β-amino acids reduce ternary complex stability and alter the translation elongation mechanism

**DOI:** 10.1101/2024.02.24.581891

**Authors:** F. Aaron Cruz-Navarrete, Wezley C. Griffin, Yuk-Cheung Chan, Maxwell I. Martin, Jose L. Alejo, S. Kundhavai Natchiar, Isaac J. Knudson, Roger B. Altman, Alanna Schepartz, Scott J. Miller, Scott C. Blanchard

## Abstract

Templated synthesis of proteins containing non-natural amino acids (nnAAs) promises to vastly expand the chemical space available to biological therapeutics and materials. Existing technologies limit the identity and number of nnAAs than can be incorporated into a given protein. Addressing these bottlenecks requires deeper understanding of the mechanism of messenger RNA (mRNA) templated protein synthesis and how this mechanism is perturbed by nnAAs. Here we examine the impact of both monomer backbone and side chain on formation and ribosome-utilization of the central protein synthesis substate: the ternary complex of native, aminoacylated transfer RNA (aa-tRNA), thermally unstable elongation factor (EF-Tu), and GTP. By performing ensemble and single-molecule fluorescence resonance energy transfer (FRET) measurements, we reveal the dramatic effect of monomer backbone on ternary complex formation and protein synthesis. Both the (R) and (S)-β^2^ isomers of Phe disrupt ternary complex formation to levels below *in vitro* detection limits, while (R)- and (S)-β^3^-Phe reduce ternary complex stability by approximately one order of magnitude. Consistent with these findings, (R)- and (S)-β^2^-Phe-charged tRNAs were not utilized by the ribosome, while (R)- and (S)-β^3^-Phe stereoisomers were utilized inefficiently. The reduced affinities of both species for EF-Tu ostensibly bypassed the proofreading stage of mRNA decoding. (R)-β^3^-Phe but not (S)-β^3^-Phe also exhibited order of magnitude defects in the rate of substrate translocation after mRNA decoding, in line with defects in peptide bond formation that have been observed for D-α-Phe. We conclude from these findings that non-natural amino acids can negatively impact the translation mechanism on multiple fronts and that the bottlenecks for improvement must include consideration of the efficiency and stability of ternary complex formation.

## INTRODUCTION

The mechanism of ribosome-catalyzed polypeptide polymerization offers the opportunity to perform template-driven synthesis of non-protein polymers with benefits for both fundamental research and biomedicine. Over the last several decades, substantial progress has been made towards expanding the genetic code to expand the expressed proteome well beyond the 20 natural α-amino acids. More than 200 different non-natural α-amino acids (nnAA) and a handful of α-hydroxy acids can be incorporated into proteins^1–5^. There are now several examples of β^2^- and/or β^3^-monomers that have been introduced into proteins in cells, either directly^6–8^ or via rearrangement^9^. Robust methods of α- and non-α nnAA incorporation into proteins is expected to promote the development of new tools to probe structure-function relationships, discover catalysts, and advance therapeutic approaches.

The most widely adopted nnAA-incorporation strategies require the transplantation of translation components from orthogonal biological systems into model organisms (e.g., *Escherichia coli*) to promote selective formation of the desired aminoacyl-tRNA^10,11^. Inefficient suppressor tRNA aminoacylation (charging) initially represented a severe bottleneck, which was largely overcome through the engineering of native aminoacyl tRNA synthases (aaRS)^12,13^. Such strides have permitted the incorporation of α-amino acids with distinct non-natural side chains into otherwise native proteins^11,14^. However, the translational machinery did not evolve to support the translation of components with alternative backbones, and significant evidence exists that multiple kinetic bottlenecks likely exist^7,15,16^. Here we identify kinetic bottlenecks that exist during the delivery and accommodation of acylated tRNA by the ribosome.

During translation, aa-tRNAs are delivered to the ribosome in ternary complex with EF-Tu (eEF1A in eukaryotes) and GTP. The three domains (DI-DIII) of EF-Tu directly engage the amino acid backbone and side chain as well as the TψC and acceptor stems of tRNA to form a high-affinity (ca. 10-100 nM) complex (**Fig.1A**)^17–19^. EF-Tu exhibits distinct affinities for each aa-tRNA species ^20,21^. The prevailing hypothesis is that this variance ensures uniform decoding speeds by balancing the spring-like forces that accumulate in aa-tRNA during initial selection and proofreading steps of tRNA selection on the ribosome that ultimately lead to aa-tRNA dissociation from EF-Tu, allowing the aminoacylated 3’-CCA terminus of the tRNA to enter the ribosome’s peptidyltransferase center^22–24^. During proofreading selection, which occurs after GTP hydrolysis at the end of initial selection, rate-limiting conformational changes in EF-Tu responsible for triggering its dissociation from aa-tRNA and the ribosome contribute to substrate discrimination by allowing additional time for near- and non-cognate aa-tRNAs to dissociate^22,25–30^. Together, initial selection and proofreading selection ensure an error rate of approximately one in 1,000-10,000 mRNA codons for natural α-amino acids^31–33^.

**Figure 1.**
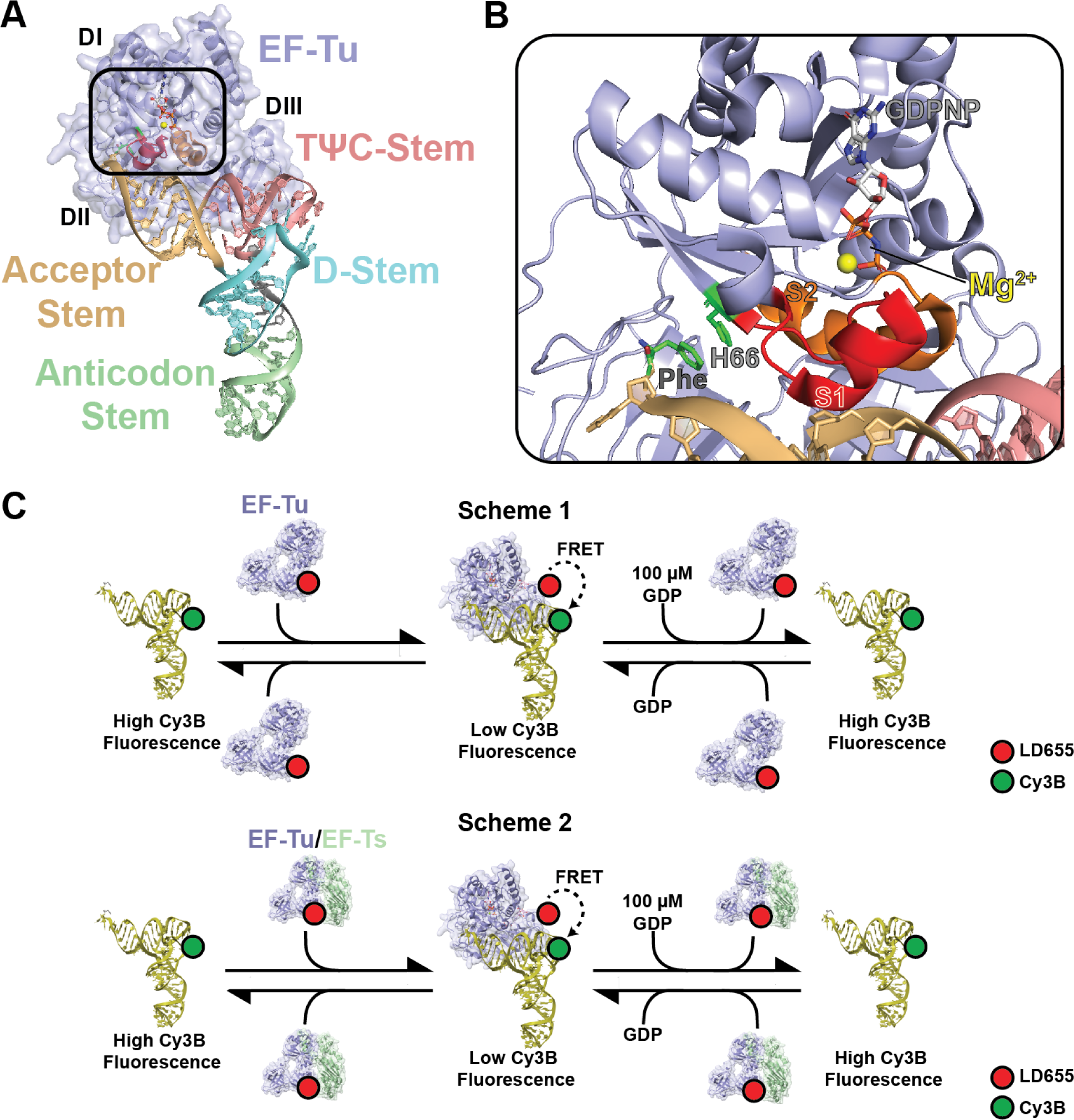
Structures of ternary complex and EF-Tu/Ts. **A)** Crystal structure of ternary complex (PDB: 1OB2) highlighting the different structural domains of tRNA (variegated) and EF-Tu (blue). The box signifies the zoom of the amino acid binding pocket of EF-Tu that is shown in panel B. **B)** Zoomed image of the EF-Tu amino acid binding pocket from the boxed region in panel A highlighting specific functional elements for ternary complex stability. The π-π stacking interaction between phenylalanine (Phe) aminoacylated to the 3’ end of tRNA^Phe^ (yellow) and histidine 66 (H66) in EF-Tu. The switch 1 (S1) and switch 2 (S2) helices which coordinate to the GTP in the nucleotide binding pocket are in red and orange, respectively. The coordinated magnesium (Mg^2+^) is shown in yellow and the non-hydorlyzable GDPNP analog is shown adjacent to the magnesium in stick form. **C)** Cartoon schematics of the FRET-based ensemble ternary complex formation assay, with LD655-labeled EF-Tu (blue) without (Scheme 1) or with (Scheme 2) EF-Ts (green) (400 nM) is injected into a cuvette containing 5 nM Cy3B-labeled tRNA^Phe^ (wheat). Formation of ternary complex results in rapid quenching of Cy3B fluorescence via FRET that can be recovered upon dissociation after injection of excess.

The active (GTP-bound) form of EF-Tu harbors a precisely formed binding pocket for the α-amino acid backbone as well as a relatively spacious cavity for the side chains of the 20 naturally occurring amino acids. Upon aa-tRNA engagement, all three EF-Tu domains (DI-III) collapse around the CCA-3’ end of the tRNA acceptor stem to position the constituent components of the amino acid side chain, which concomitantly restructures the switch 1 helix (S1) to engage the γ-phosphate of GTP via Mg^2+^ coordination to yield the thermodynamically stable ternary complex (**Fig. 1B**)^34,35^. This stability strictly depends on tRNA aminoacylation and the presence of a properly positioned γ-phosphate moiety^19,36^. In line with this exquisite sensitivity, *in vitro* transcribed tRNAs mis-acylated with native amino acids can exhibit binding affinities for EF-Tu(GTP) that are reduced by up to ∼5000-fold^20^, which affect both the speed and fidelity of tRNA selection on the ribosome^37–39^. However, the precise contributions of the amino acid backbone to EF-Tu(GTP) affinity have yet to be fully explored.

In bacteria, ternary complex formation is catalyzed by the conserved, EF-Tu-specific guanosine nucleotide exchange factor (GEF), EF-Ts. Under nutrient-rich conditions where GTP concentration far exceeds that of GDP, EF-Ts ensures rapid and abundant ternary complex formation so that ternary complex formation is not rate-limiting to protein synthesis^19,36^. Under nutrient-poor conditions when GDP concentrations are elevated, EF-Ts instead facilitates ternary complex disassembly, reducing protein synthesis and other energy-intensive cellular processes^19,36^. The impact of EF-Ts on both ternary complex formation and dissociation indicates that nucleotide exchange and S1 restructuring are dynamic processes that can be influenced by EF-Ts both in the absence or presence of aa-tRNA.

Here we use ensemble and single-molecule FRET-based kinetic assays, together with kinetic simulations, to investigate the effects of both non-natural α-amino acids (specifically, those with a non-natural side chain and a natural α-backbone) and non-natural backbones (specifically, those with a natural side chain and a non-natural β^2^-or β^3^-amino acid backbone) on the kinetics and thermodynamics of ternary complex formation and tRNA elongation on the ribosome^19,36^. Consistent with its widespread use by diverse research labs, our investigations show that the metrics of both ternary complex formation and tRNA selection are virtually identical for tRNAs acylated with α-Phe or the non-natural α-amino acid *para*-azido-phenylalanine (*p*-Az-Phe). By contrast, the kinetics of both ternary complex formation and tRNA selection were altered for backbone-modified monomers. The presence of either (R) or (S)-β^2^ Phe in place of L-α-Phe disrupts ternary complex formation to levels below our *in vitro* detection limits, whereas (R)- and (S)-β^3^-Phe reduce ternary complex stability by approximately an order of magnitude. These deficiencies are exacerbated by thermally stable elongation factor EF-Ts, the cellular guanosine nucleotide exchange factor for EF-Tu, and by mutations in both EF-Tu and tRNA previously reported to stabilize ternary complex using *in vitro* transcribed tRNAs^40,41^. In line with these observations, ribosomes fail to recognize (R)- and (S)-β^2^-Phe-charged tRNAs as substrates, while the utilization of (R)- and (S)-β^3^-Phe stereoisomers was significantly impaired relative to native L-α-Phe. The reduced EF-Tu affinities of tRNAs acylated with either (R)-or (S)-β^3^-Phe also precipitated defects in the mRNA decoding mechanism, where the proofreading stage of the tRNA selection process immediately after GTP hydrolysis was ostensibly bypassed. Following incorporation into the ribosome, tRNAs charged with (R)- and (S)-β^3^-Phe stereoisomers were both competent for translocation. However, as predicted using recent metadynamics simulations^16^, (R)-β^3^-Phe appeared to exhibit order of magnitude defects in the rate of peptide bond formation, which dramatically reduced the rate of substrate translocation from hybrid-state tRNA positions. We conclude from these findings that the efficiency of ternary complex formation and its thermodynamic stability are key determinants of nnAA incorporation into proteins and that engineering opportunities exist to enable their more efficient utilization.

## RESULTS

### Quantifying ternary complex formation

To directly investigate how various types of non-natural amino acids impact the kinetic features of ternary complex formation, we employed an ensemble approach in which the rate and extent of ternary complex formation was tracked via FRET^19,36^. In this assay, EF-Tu carrying an LD655 fluorophore at its C-terminus quenches the fluorescence of a Cy3B fluorophore attached to the 3-amino-3-carboxypropyl (acp^3^) modification at position U47 of native tRNA^Phe^ when the ternary complex forms (**Fig. 1C**). Reactions were initiated upon stopped-flow addition of 400 nM EF-Tu-LD655 to a solution of 5 nM L-α-Phe-tRNA^Phe^-Cy3B in a 1.2 mL cuvette with stirring at 25 °C (**Methods**). Under these conditions, the observed pseudo first-order, apparent rate constant (*k*_obs_) for Cy3B fluorescence quenching was 0.12 ± 0.002 s^-1^, reaching ∼65% quenching at equilibrium (**Fig. 2A**; **Table 1**). The observed fluorescence quenching amplitude agrees with the close proximity of Cy3B and LD655 fluorophores within the ternary complex (**Fig. 1**)^19^. Consistent with the GTP requirement for ternary complex formation, rapid stopped-flow addition of 100 µM GDP to the reaction restored Cy3B fluorescence to ∼80% of the initial intensity with an apparent rate constant (*k*_off_) of 0.005 ± 0.001 s^-1^ (**Fig. 2A**; **Table 1**). Identical experiments performed using deacylated tRNA^Phe^-Cy3B in place of L-α-Phe-tRNA^Phe^-Cy3B resulted in no Cy3B quenching (**Fig. 2A**), congruous with the selectivity of EF-Tu for aminoacylated tRNA.

**Figure 2.**
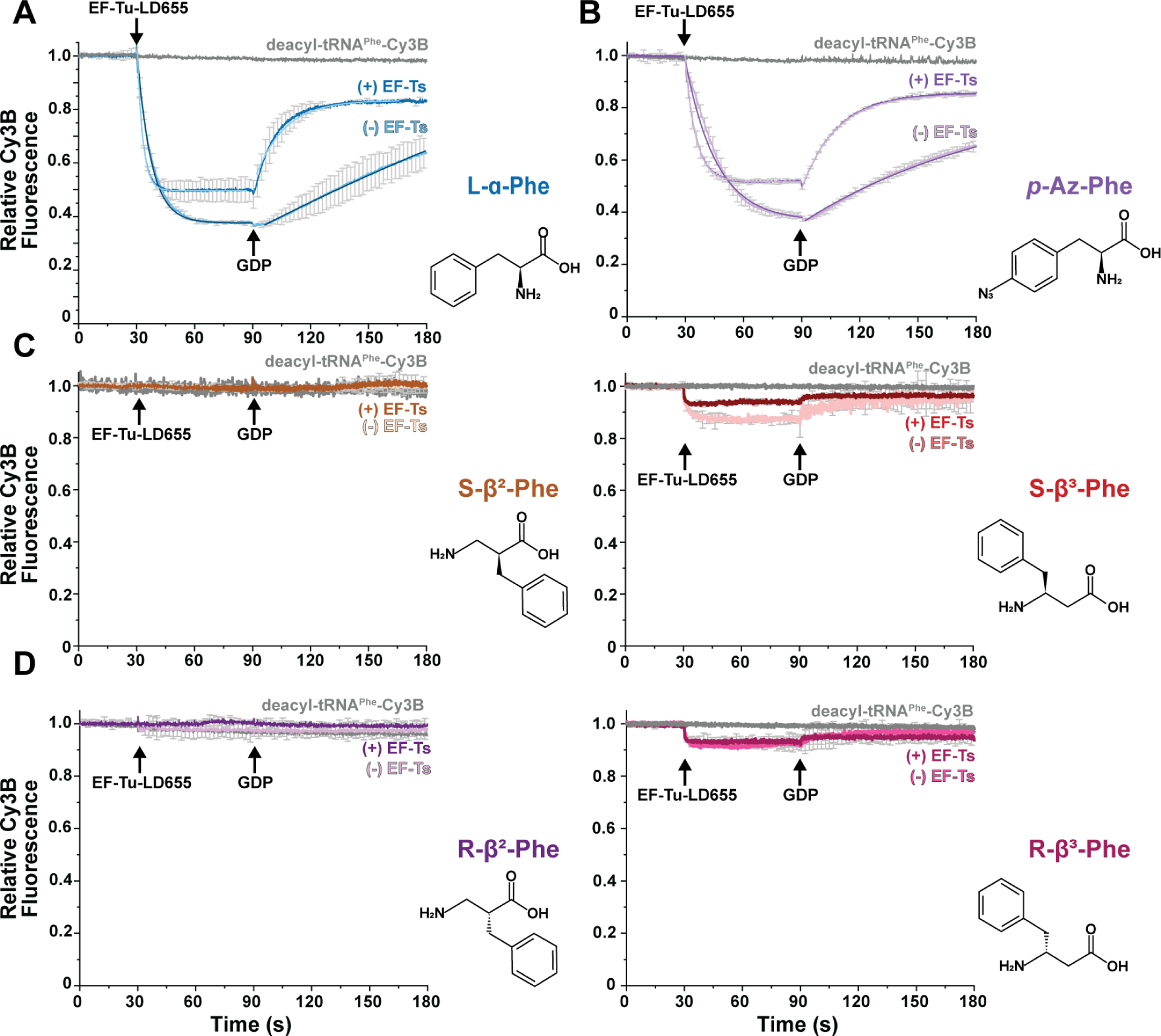
Non-natural amino acids severely disrupt ternary complex stability. Ensemble ternary complex assays tracking the apparent rates (*k_obs_*) of Cy3B relative fluorescence changes upon mixing with LD655-labeled EF-Tu (-/+) EF-Ts (400 nM) in the presence of GTP (100 µM GDP) with Cy3B-labeled tRNA^Phe^ (5 nM) charged with specific amino acid monomers with **A)** L-α-Phe. **B)** *p*-azido-Phe (pAzF). **C)** S-β^2^-Phe (left); S-β^3^-Phe (right). **D)** R-β^2^-Phe (left); R-β^3^-Phe (right). Structures of amino acid monomers are shown to the right of each plot. Error bars represent standard deviation from two experimental replicates.

**Table 1.**
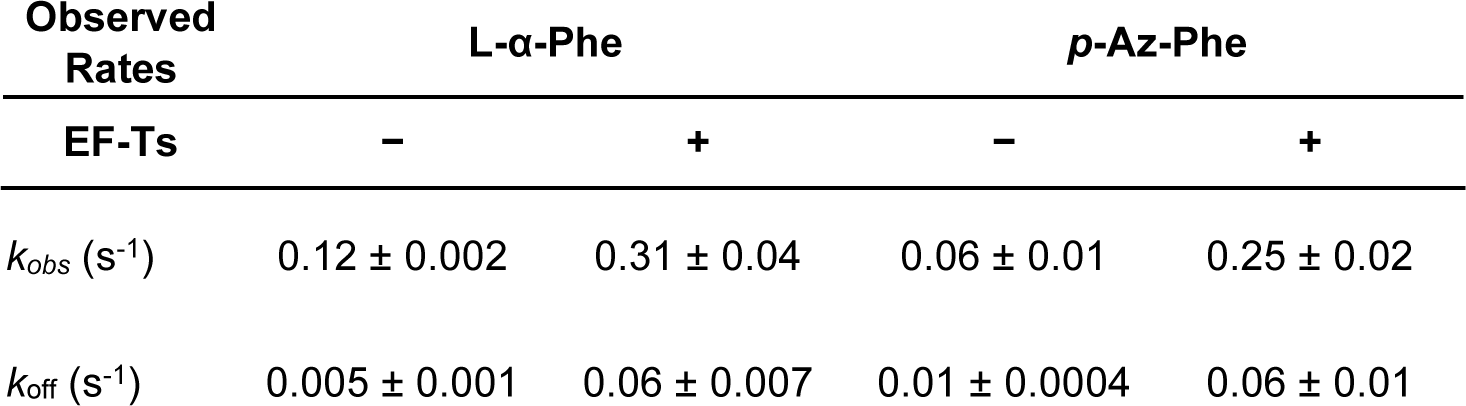
Apparent reaction rates estimated by single-exponential fitting of the ensemble ternary complex formation experiments shown in Figure 2. Uncertainty estimates represent S.D. from two experimental replicates. Single-exponential fits exhibited R^2^ values >0.95. The L-α-Phe rates compare well with previously reported results ^36^.

When analogous experiments were performed with the nucleotide exchange factor EF-Ts present in a 1:1 ratio relative to EF-Tu, *k_obs_* increased to 0.31 ± 0.04 s^-1^ and the extent of Cy3B fluorescence quenching at equilibrium was reduced to ∼50%. Addition of 100 µM GDP to the same reaction restored the Cy3B fluorescence amplitude with a *k*_off_ that was approximately 10-fold faster than with EF-Tu alone (0.06 ± 0.007 s^-1^). As for EF-Tu alone, the fluorescence intensity prior to ternary complex formation was nearly fully restored, although not to the full extent due to residual ternary complex formation under equilibrium conditions (**Fig 1C**, **scheme 2**; **Fig. 2A**). These observations are consistent with previously reported affinities of EF-Tu(GTP) for aa-tRNA (ca. 10-100 nM) and prior conclusions that EF-Ts can engage with EF-Tu in ternary complex to catalyze nucleotide exchange, thereby accelerating ternary complex dissociation rates after GDP addition by ∼20-fold^19,36^.

### Ternary complex stability is disrupted by nnAAs

With this foundation, we next asked how the rates of ternary complex formation and stability were affected when tRNA^Phe^-Cy3B was aminoacylated with non-natural amino acids such as *p*-Az-Phe, (R)- and (S)-β^2^-Phe and (R)- and (S)-β^3^-Phe. These nnAA were used to acylate native tRNA^Phe^-Cy3B using flexizyme^42–44^. Each charged species was purified by FPLC, flash frozen in aliquots, and verified as >95% intact prior to use (**Methods; Fig. S1A, B**). We confirmed that flexizyme-charged L-α-Phe-tRNA^Phe^-Cy3B exhibited nearly identical formation/dissociation kinetics and fluorescence quenching/recovery amplitudes as the enzymatically charged species (**Fig. S2**), validating this experimental tRNA aminoacylation strategy for kinetic assays^6,15,43,45^.

Using experimental conditions identical to those described above, we next examined the kinetics of ternary complex assembly usingtRNA^Phe^-Cy3B that was acylated with *p*-Az-Phe, a nnAA successfully incorporated into proteins by multiple laboratories^10,13,46,47^. Stopped-flow addition of EF-Tu-LD655 to *p-*Az-Phe-tRNA^Phe^-Cy3B resulted in Cy3B fluorescence quenching characterized by a *k*_obs_ of 0.06 ± 0.01 s^-1^; the extent of fluorescence quenching reached ∼65% at equilibrium (**Fig. 2B**; **Table 1**). As observed for the complex of L-α-Phe-tRNA^Phe^-Cy3B, rapid addition of GDP slowly restored Cy3B fluorescence; in this case, the measured *k*_off_ was 0.01 ± 0.0004 s^-1^ (**Fig 2B** and **Table 1**). In presence of EF-Ts, ternary complex formation again proceeded more rapidly (*k*_obs_ = 0.25 ± 0.02 s^-1^) and the fluorescence quenching reached an amplitude of ∼50% at equilibrium (**Fig 2B** and **Table 1**). Ternary complex dissociation upon addition of GDP proceeded with a *k*_obs_ of 0.06 ± 0.02 s^-1^, restoring Cy3B fluorescence to a ∼80% of the initial intensity. Overall, the kinetics parameters measured for assembly and disassembly of ternary complexes containing *p-*Az-Phe-tRNA^Phe^-Cy3B were similar to those measured for complexes containing L-α-Phe-tRNA^Phe^-Cy3B, Thus, the presence of the *p*-N_3_ side chain on Phe has, as expected, only a modest impact on ternary complex formation and stability. These results are entirely consistent with the functionalized phenyl sidechain being readily accommodated by EF-Tu.

We next investigated the kinetics of ternary complex assembly and disassembly when tRNA^Phe^-Cy3B was aminoacylated with the (R)-or (S)-enantiomers of β^2^-or β^3^-Phe. Many β^2^- and β^3^-amino acids have been introduced into short peptides using small scale *in vitro* translation reactions^48,49^ and a few β^3^-amino acids have been introduced into proteins in cell lysates^50,51^. Moreover, one β^3^-Phe derivative^6^, three β^3^-aryl derivatives^8^, and one β^2^-hydroxy acid^7^ have been introduced into proteins in cells. Yet in none of these cases was the level of incorporation especially high, perhaps because of impaired delivery to the ribosome by EF-Tu^52,53^.

Thus, using experimental conditions identical to those described above, we examined ternary complex formation using tRNA^Phe^-Cy3B that was acylated with the (R)- and (S)-enantiomers of β^2^- and β^3^-Phe (**Methods; Fig. 2**). Under these conditions we detected no evidence for ternary complex formation in reactions containing (R)-or (S)-β^2^-Phe-tRNA^Phe^-Cy3B (**Fig**. **2C**), while reactions containing (R)-or (S)-β^3^-Phe-tRNA^Phe^-Cy3B exhibited detectable, yet greatly reduced levels of Cy3B fluorescence quenching (∼10-15% vs. ∼60-65% with L-α-Phe). Quenching was reversed upon addition of 100 µM GDP, as expected for quenching that resulted from ternary complex formation (**Fig**. **2C** and **2D**). Also consistent with ternary complex formation was the observation that addition of EF-Ts to the ternary complex assembly reactions containing (R)-or (S)-β^3^-Phe-tRNA^Phe^-Cy3B attenuated Cy3B fluorescence quenching. Due to the impaired signal amplitudes of these reactions, reliable *k_obs_* and *k*_off_ rates could not, however, be estimated.

### β^2^-Phe and β^3^-Phe monomers negatively impact the kinetic features of ternary complex formation

We next employed a micro-volume stopped-flow system (µSFM, Biologic) (**Methods**) to perform a thorough kinetic study of ternary complex formation for tRNAs carrying (R) or (S)-β^3^-Phe, which showed the highest fluorescence quenching of the four acylated tRNAs studied above. Using this approach, we measured bimolecular association rate constants for EF-Tu binding to tRNA^Phe^-Cy3B acylated with either L-α-Phe-or (S)-β^3^-Phe-charged tRNA^Phe^ by tracking how *k*_obs_ changed as a function of the concentration of EF-Tu or EF-Tu/Ts. Upon rapid mixing of 5 nM L-α-Phe-charged tRNA^Phe^-Cy3B with a large excess of EF-Tu (0.2 – 2 µM) and GTP (1 mM), we observed a linear increase in *k_obs_* as a function of both [EF-Tu] and [EF-Tu/Ts], consistent with pseudo-first order binding kinetics (**Figure 3A** and **3B**). In line with previous literature^19,36^, in the absence of EF-Ts, the bimolecular rate constant (*k*_on_) was 5.0 ± 0.1 µM^-1^ s^-1^ and the dissociation rate constant (*k*_off_) was 0.2 ± 0.1 s^-1^ (**Table 2**). With equimolar EF-Tu/Ts, *k*_on_ and *k*_off_ increased by 1.6-and 4-fold, respectively^19,36^. Consistent with our initial findings, when L-α-Phe was substituted with (S)-β^3^-Phe, we observed both a lower *k*_on_ (1.0 ± 0.1 µM^-1^ s^-1^) and a higher *k*_off_ (1 ± 0.1 s^-1^) (**Table 2**). In the presence of EF-Ts, *k*_on_ and *k*_off_ increased by 3-fold and 1.8-fold, respectively (**Table 2**). For (R)-β^3^-Phe, the fluorescence intensity changes were still too small for reliable estimations of its kinetic parameters (**Fig. S3**). Additionally, we were unable to rescue the observed defects using tRNA (C49A, G65U) and EF-Tu (N273A) mutations previously reported to stabilize ternary complex formation (**Methods; Fig. S3**).

**Figure 3.**
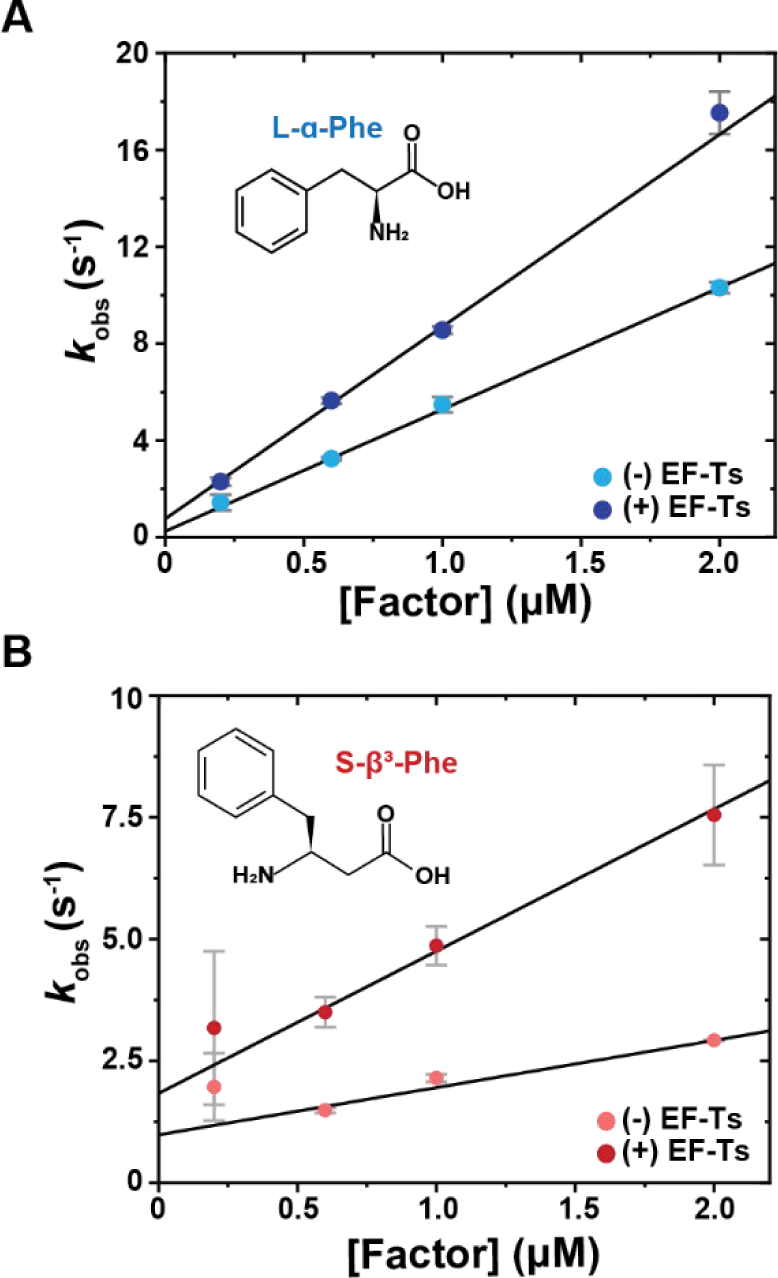
Rapid, pre-steady state measurements of ternary complex formation rate constants experiments. **A)** Plot showing the apparent rate (*k*_obs_) of ternary complex formation for Cy3B-labeled L-α-Phe-tRNA^Phe^ as a function of EF-Tu (sky blue) and equimolar EF-Tu/Ts (navy blue) concentrations performed in the presence of 100 µM GTP. **B)** Analogous plots of *k*_obs_ for Cy3B-labeled S-β^3^-Phe-tRNA^Phe^ in the absence (salmon) and presence (red) of equimolar EF-Ts. **C)** Kinetic framework for ternary complex formation based on prior literature ^19,36,79^, indicating ternary complex formation-inactive (grey) and active (black) species of EF-Tu. Error bars represent S.D. from 3-7 experimental replicates.

**Table 2.** Kinetic and thermodynamic parameters for aa-tRNA^Phe^ binding to EF-Tu-LD655 in the presence and absence of EF-Ts from the experiments in Figure 3. Asterisks (*) indicate rate and equilibrium constants derived from experiments in the presence of EF-Ts. K_D_ values were estimated from the ratio K_D_ = *k*_off_ /*k*_on_ determined from titration experiments in Figure 3. Free energies were calculated using the relationship Δ*G°* = *–RT*ln(1/KD) ^20^

| Rate<br>constant | L-α-Phe | | S-$\beta^3$-Phe | |
| --- | --- | --- | --- | --- |
| EF-Ts | – | + | – | + |
| $k_{\text{on}}$ ($\mu\text{M}^{-1} \text{ s}^{-1}$) | $5.0 \pm 0.1$ | $8.0 \pm 0.3^*$ | $1.0 \pm 0.1$ | $3.0 \pm 0.3^*$ |
| $k_{\text{off}}$ ($\text{s}^{-1}$) | $0.2 \pm 0.1$ | $0.8 \pm 0.2^*$ | $1.0 \pm 0.1$ | $1.8 \pm 0.2^*$ |
| $K_{\text{D}}$ (est.) | 40 nM | 100 nM* | 1 $\mu\text{M}$ | 0.6 $\mu\text{M}^*$ |
| $\Delta G^\circ$<br>(kcal/mol) | -11.5 | -9.6 | -4.1 | -4.4 |

From these findings we estimated equilibrium dissociation constants (*K*_D_) for the ternary complexes of EF-Tu with L-α-Phe-tRNA^Phe^-Cy3B of approximately 40 and 100 nM in the absence and presence of EF-Ts, respectively (**Table 2**). Similar analyses estimated ternary complex *K*_D_ values of ∼1000 and ∼600 nM for the analogous complexes containing (S)-β^3^-Phe, in the absence and presence of EF-Ts, respectively (**Table 2**). From these *K*_D_ estimates we calculated Δ*G*° values of −9 to −11 kcal/mol for complexes containing L-α-Phe, consistent with previous literature^20,41,54^, while the Δ*G*° for complexes containing (S)-β^3^-Phe were only −4.1 to −4.4 kcal/mol. Hence, the stability of the (S)-β^3^-Phe-containing ternary complex would be outside of the thermodynamic range for ternary complexes that support efficient translation, as both tightly and loosely bound aa-tRNAs to EF-Tu impair translation by slowing peptide bond formation^55^ or aa-tRNA delivery to the ribosome, respectively^54^ (**Table 2**).

These data support the hypothesis that both β^2^- and β^3^-Phe monomers interfere with EF-Tu engagement by both reducing the efficiency with which EF-Tu productively engages acyl-tRNA as well as the overall stability of the ternary complex. Despite an approximate 25-fold reduction in affinity for (S)-β^3^-Phe-tRNA^Phe^, kinetic simulations based on a simplified framework (**Fig. S4A**), suggest that (S)-β^3^-Phe-ternary complexes can be populated significantly in a cellular context, where EF-Tu and EF-Ts are present at ∼µM concentrations (**Fig. S4B and S4C**)^56^. We speculate that increasing cellular EF-Tu concentrations may only partially alleviate incorporation deficiencies due to complex instability.

### β-Phe-charged tRNAs are poorly accommodated on the ribosome and ostensibly bypass proofreading steps

To examine how ternary complexes formed with nnAAs are incorporated by the ribosome, we employed smFRET imaging methods to directly monitor both the frequency of productive ternary complex binding to the ribosome as well as the process by which an aa-tRNA is released from EF-Tu and incorporated into the ribosomal aminoacyl (A) site^22,24,29^. This established approach tracks FRET between a donor fluorophore linked to a peptidyl-tRNA bound within the ribosomal P-site and an acceptor fluorophore linked to the incoming aa-tRNA (**Fig. 4A**).

**Figure 4.**
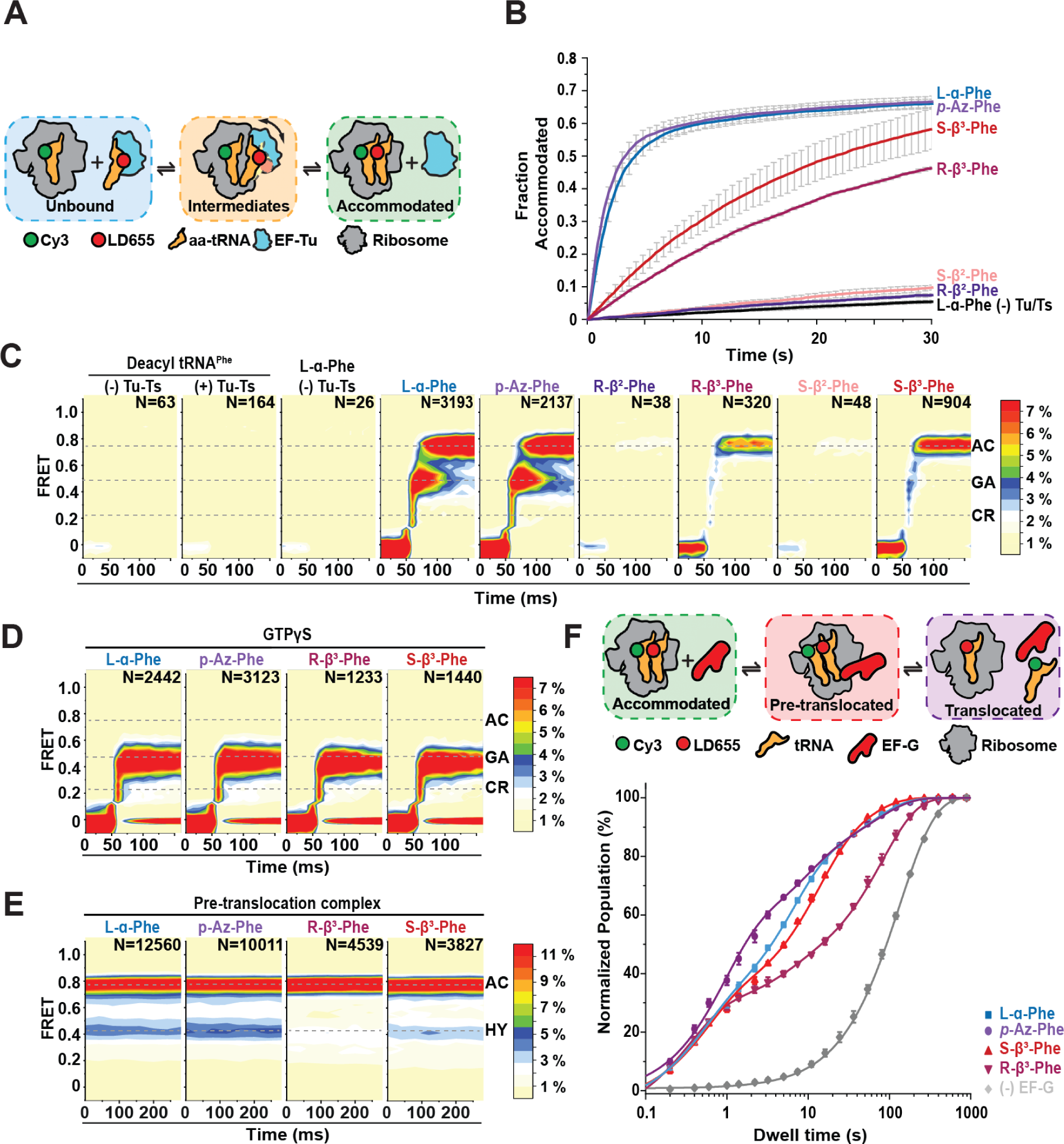
nnAA-tRNA selection on the bacterial ribosome studied by smFRET. **A)** Schematic describing the smFRET tRNA selection experimental design, where LD655-labeled aa-tRNA^Phe^ is delivered to surface-tethered ribosomes (blue) bearing Cy3-fMet-tRNA^fMet^ in the P site and tRNA selection intermediates (tan), and the fully accommodated tRNA (green), are distinguished based on their distinct FRET values. **B)** Fraction of ribosomes bearing a fully accommodated aa-tRNA as a function of time (**Methods**). Data were collected at 100 ms time resolution; experiments were performed in triplicate. **C-D)** 2D histograms of smFRET traces containing productive accommodation events with GTP **(C)** or stalled events with GTPγS **(D)** (see methods) (N, number selected), post-synchronized to the appearance of FRET above baseline, with the specified amino acids. Contour plots were normalized (scale at right) to the L-α-Phe positive control. Dashed lines indicate FRET efficiency values of tRNA selection intermediates used in kinetic modeling. Data were collected at 10 ms time resolution. **E)** 2D histograms of smFRET traces of accommodated aa-tRNA after buffer exchange and ∼5 min equilibration time, showing both classical (accommodated) and hybrid (P/E, A/P) tRNA conformations. Data were collected at 40 ms time resolution. **F)** Schematic and fraction of productive elongation cycles over time from tRNA selection and translocation experiments in presence of 10 µM EF-G. Line represent fits to one (L-α-Phe without EF-G) or three (L-α-Phe, *p*-Az-Phe, (R)- and (S)-β^3^-Phe) exponential distributions. Three different dwell time regimes are observed: the hybrid state (∼0.5 s), the classical state (∼8 s) and the “slow” state (∼60 s). Data were collected at 200 ms time resolution. For all tRNA selection experiments, LD655-labeled aa-tRNA^Phe^ and EF-Tu/Ts concentrations were 12.5 nM and 125 nM, respectively. For all translocation experiments, LD655-labeled aa-tRNA^Phe^ and EF-Tu/Ts concentrations were 25 nM and 250 nM, respectively.

To perform these studies, bacterial ribosome complexes were programmed with a synthetic 5’-biotinylated mRNA that positions Cy3-fMet-tRNA^fMet^ in the P site and a UUC codon in the A site. Initiation complexes were then tethered via a biotin-streptavidin bridge proximal to an optically transparent, polyethylene glycol (PEG)-passivated surface. Single-molecule tRNA selection experiments were subsequently initiated by stopped-flow injection of ternary complexes formed with flexizyme-charged tRNA^Phe^-LD655, where EF-Tu/Ts (125 nM) is in 10-fold excess over aa-tRNA (12.5 nM) (**Methods**).

We first assessed the apparent rates and extents of ternary complex utilization by the ribosome by collecting movies at low-time resolution (100 ms per video frame) where photobleaching is minimized. Consistent with our ensemble ternary complex formation studies (**Figs. 2, 3**), *p*-Az-Phe-tRNA^Phe^-LD655 was utilized as efficiently as L-α-Phe-tRNA^Phe^-LD655 (**Fig. 4B**; **Table 3**). Both (R)- and (S)-β^2^-Phe-LD655 were not utilized by the ribosome, while both (R)- and (S)-β^3^-Phe-tRNA^Phe^-LD655 were utilized >10-fold less compared to L-α-Phe-tRNA^Phe^-LD655, defects that we attribute to defects in ternary complex formation (**Fig. 4B**; **Table 3**).

**Table 3.** Apparent aa-tRNA accommodation rates on the ribosome calculated from the experiments in Figure 4. Uncertainty estimates represent S.D. from three experimental replicates.

| Amino acid | $k_{1\text{ obs}} (\text{s}^{-1})$ | $k_{2\text{ obs}} (\text{s}^{-1})$ |
| --- | --- | --- |
| L-α-Phe (-) Tu-Ts | $0.025 \pm 0.002$ | N/A |
| L-α-Phe | $0.46 \pm 0.04$ | $0.033 \pm 0.001$ |
| *p*-Az-Phe | $0.59 \pm 0.03$ | $0.036 \pm 0.002$ |
| R-$\beta^2$-Phe | $0.024 \pm 0.003$ | N/A |
| R-$\beta^3$-Phe | $0.041 \pm 0.002$ | N/A |
| S-$\beta^2$-Phe | $0.024 \pm 0.001$ | N/A |
| S-$\beta^3$-Phe | $0.056 \pm 0.006$ | N/A |

To gain more specific insights into the tRNA selection mechanism after ternary complex binding to the ribosome, we performed analogous investigations at 10-fold higher time resolution (10 ms per video frame). As previous smFRET studies of bacterial tRNA selection have shown^22,29^, experiments of this kind allow direct assessment of codon recognition (CR), GTPase activation (GA), and accommodation (AC) as each reaction endpoint exhibits a distinct FRET efficiency value (∼0.21, 0.45, and 0.75, respectively; **Methods**).

By computationally isolating FRET trajectories reflecting productive ternary complex binding events to individual ribosomes (**Methods**)^22^ (i.e. those that stably accommodate) (**Fig. 4C**), we found that L-α-Phe and *p-*A*z*-Phe exhibited similar progression probabilities through tRNA selection, including the rate-limiting progression from the GTPase-activated state (∼0.45, GA) to the fully accommodated state (∼0.75, AC) during proofreading selection. As expected, tRNA selection events were ostensibly not observed for ternary complexes in which tRNA was acylated with (R)-or (S)-β^2^-Phe, whereas we observed ∼5-10-fold fewer productive tRNA selection events for (R)- and (S)-β^3^-Phe ternary complexes (**Fig. 4C**). Strikingly, these rare events of (R)- and (S)-β^3^-Phe tRNA accommodation exhibited much more rapid passage of the intermediate states of tRNA selection compared to L-α-Phe.

To test whether the tRNA selection process for (R)- and (S)-β^3^-Phe bypassed GTP hydrolysis, we performed identical smFRET experiments with the non-hydrolyzable GTPγS analog to stall tRNA selection at the GTP hydrolysis step at the end of initial selection^36^. In the presence of GTPγS, both (R)- and (S)-β^3^-Phe-tRNA^Phe^-LD655 were efficiently stalled in the GA state, as were both L-α- and *p*-Az-Phe-tRNA^Phe^-LD655 (**Fig. 4D**). These findings reveal that GTP hydrolysis is indeed required for (R)- and (S)-β^3^-Phe-tRNA^Phe^ to for GA state passage and the proofreading process. The observation that proofreading is significantly more rapid for (R)- and (S)-β^3^-Phe-tRNA^Phe^ supports the notion that conformational changes in EF-Tu during proofreading related to 3’-CCA release are rate-limiting to the tRNA selection process^22,29^ and that this impact can be specifically attributed to the β-Phe monomers. Reduced thermodynamic stability leads to more rapid EF-Tu dissociation from aa-tRNA and likely the ribosome.

Since we were able to observe some β^3^-Phe accommodation on the ribosome, we next wanted to assess if (R)- and (S)-β^3^-Phe-tRNA^Phe^ impact later steps of the elongation cycle, including peptide bond formation and EF-G catalyzed translocation. To examine these steps, first we examined the equilibrium between classical hybrid state pre-translocation complex conformations (A/P, P/E and A/A, P/E) that are adopted spontaneously after tRNA selection^57–59^ (**Fig. 4E**). L-α-Phe and *p*-Az-Phe showed a 64:36 (± 2) split ratio between classical and hybrid pre-translocation complexes. Notably, (S)-β^3^-Phe did not significantly affect the classical-hybrid split ratio (67:33 ± 3), while (R)-β^3^-Phe preferentially adopted classical conformations (74:26 ± 2; *p* < 0.05; **Methods**). These results indicate that, while (S)-β^3^-Phe behaves similarly to its α-amino acid counterparts, (R)-β^3^-Phe also perturbs spontaneous transitions to translocation-ready hybrid states^23^. Hence, incorporation of (R)-β^3^-Phe into proteins is predicted to increase the frequency of elongation pauses.

We next performed analogous smFRET experiments under cell-like conditions, where productive tRNA selection events are rapidly followed by translocation. To initiate the complete elongation cycle, we stopped-flow injected ternary complexes formed with LD655-labeled tRNA^Phe^ (250nM EF-Tu/Ts, 25nM aa-tRNA) together with EF-G (10 µM) in the presence of 1 mM GTP. In this experiment, the appearance of FRET reports on aa-tRNA incorporating into the ribosome. The loss of FRET reports on dissociation of Cy3-labeled initiator tRNA^fMet^ from the E site during, or after, the process of translocation^60^ (**Fig. 4F**). Correspondingly, FRET lifetime in these experiments reports on the total duration of the elongation cycle. We measured complete elongation cycle times using ternary complexes formed with L-α-Phe, *p*-Az-Phe as well as (R)- and (S)-β^3^-Phe-tRNA^Phe^, while including a control study lacking EF-G to estimate the contribution of fluorophore photobleaching (**Fig. 4F**). Consistent with each complex undergoing EF-G catalyzed translocation, the average FRET lifetimes were substantially shorter (ca. 2.5-8 fold) than photobleaching (165.8 ± 2.8 s). In line with prior investigations of translocation^23,60–64^, the FRET lifetime distributions for each amino acid type displayed multi-modal behaviors, characterized by a very fast (∼0.3 s), fast (∼5-10 s) and slow (∼30-40 s) kinetic populations (**Table 4**). We attribute these sub-populations to pre-translocation complexes that undergo rapid translocation from hybrid state conformations, classical pre-translocation complexes that must wait for hybrid state conformations to spontaneously occur and pre-translocation (“slow”) complexes that either exhibit more substantial delays in translocation or retain deacylated tRNA in the E site after translocation is complete. L-α-Phe, *p*-Az-Phe as well as (R) and (S)-β^3^-Phe each exhibit rapidly translocating sub-populations, consistent with rapid peptide-bond formation followed by rapid translocation from hybrid state conformations. Sub-populations were also present for all four amino acids that exhibited long-lived FRET lifetimes, consistent with relatively stable, classical ribosome conformations requiring additional time to first transition to hybrid states before being able to productively engage EF-G and then rapidly translocate^23,60,62^ (**Table 4**). We note in this context that (R)-β^3^-Phe, the monomer most dissimilar to L-α-Phe, exhibited a relatively large sub-population of pre-translocation complexes that exhibit “slow” translocation, which may reflect specific deficiencies in peptide bond formation, hybrid state formation or deacylated tRNA release from the E site.

**Table 4.** Dwell times for hybrid, classical and “slow” states from the fittings to three exponential distributions (**Methods**). Photobleaching lifetime is 165.8 ± 0.03 s.

| Amino acid | Hybrid state |  | Classical state |  | “Slow” state |  |
| --- | --- | --- | --- | --- | --- | --- |
|  | Dwell time (s) | Population (%) | Dwell time (s) | Population (%) | Dwell time (s) | Population (%) |
| L-α-Phe | $0.44 \pm 0.11$ | $30.1 \pm 3.6$ | $6.12 \pm 0.68$ | $48.2 \pm 3.1$ | $59.17 \pm 3.12$ | $21.7 \pm 1.7$ |
| *p*-Az-Phe | $0.87 \pm 0.08$ | $53.3 \pm 2.2$ | $8.45 \pm 1.13$ | $26.2 \pm 1.6$ | $67.46 \pm 3.35$ | $20.5 \pm 1.2$ |
| S-$\beta^3$-Phe | $0.53 \pm 0.07$ | $33.6 \pm 2.9$ | $11.81 \pm 0.83$ | $46.7 \pm 2.2$ | $47.56 \pm 3.76$ | $19.7 \pm 2.5$ |
| R-$\beta^3$-Phe | $0.33 \pm 0.03$ | $33.9 \pm 0.9$ | $4.67 \pm 0.80$ | $12.1 \pm 1.2$ | $80.79 \pm 3.31$ | $54.0 \pm 1.1$ |

## DISCUSSION

Efficient tRNA selection by the ribosome is paramount to the transfer of genetic information from mRNA to protein^65^. Harnessing this template-driven platform through genetic code expansion to synthesize previously unknown and useful bio-polymers promises myriad tools to advance research and medicine^11^. Achieving this goal requires continual technology development to overcome bottlenecks that limit the efficiency of non-α amino acid incorporation. These bottlenecks in principle include, but are not limited to, differences in monomer uptake and cellular stability, aminoacyl-tRNA synthetase activity and fidelity, ternary complex formation of the aa-tRNA with EF-Tu(GTP), constraints present within the ribosome PTC itself, as well as other fidelity mechanisms evolved to reduce erroneous amino acid incorporation by the ribosome^13,66–69^. Here we employ both ensemble and single-molecule kinetic assays, we show that ternary complex formation and utilization by the ribosome represent significant bottlenecks for the use of β-Phe monomers.

tRNAs acylated with (R)- and (S)-β-Phe monomers are disruptive to ternary complex formation and thus their utilization by the ribosome (**Fig. 2C**, **D**; **Fig. 4B**, **C**). tRNAs acylated with (R)- and (S)-β^2^-Phe appeared unable to form ternary complex with EF-Tu under the experimental conditions examined. The incorporation of (R)- and (S)-β^2^-Phe-tRNAs into the ribosome was below the detection limits of our tRNA selection smFRET assays. By contrast, (R)- and (S)-β^3^-Phe monomers inefficiently formed ternary complex, reducing ternary complex abundance and thus the number of detectable tRNA selection events per unit time (**Fig. 4B, C**). Strikingly, the proofreading stage of tRNA selection was accelerated for the events observed, consistent with faster rates of EF-Tu dissociation from both tRNA and the ribosome. We infer from these findings that β^3^-Phe monomers lower ternary complex stability to an extent that the proofreading stage of tRNA selection after GTP hydrolysis is ostensibly bypassed. As L-α-amino acids misacylated onto non-native tRNAs have been reported to exhibit identical accommodation rates despite evidence of ternary complex formation defects^70^, further experiments will be needed to discern whether the impact on proofreading is specific for β^3^-Phe monomers.

Our results also show that β^3^-Phe monomers exhibit significant reductions in the rate of downstream elongation reactions, including EF-G catalyzed translocation (**Fig. 4F**). (R)-β-Phe monomers, whose stereochemistry mimics that of an unnatural D-α-amino acid, showed significantly greater defects in ternary complex formation, tRNA accommodation during the proofreading stage of tRNA selection and EF-G-catalyzed translocation than their (S)-β-Phe counterparts. These observations are consistent with prior literature indicating that D-α-amino acids ((R)-α-amino acids) result in extended elongation pauses^15,45,71^. In the cell, such pauses are likely accompanied by futile cycles of EF-Tu catalyzed GTP hydrolysis^29^ and the induction of rescue pathways^72^ that may ultimately lead to compromises in cell growth and viability.

We conclude from these findings that ternary complex stability is a significant and perhaps underappreciated bottleneck that limits the incorporation of extended backbone monomers into polypeptides and proteins, both in vitro and in vivo. These observations warrant examination of the extent to which ternary complex stability and EF-Tu catalyzed tRNA selection defects impact the efficiency of nnAA utilization more broadly. Structural data^19,36,73,74^ suggest that the extended backbones of both β^2^- and β^3^-Phe may introduce steric clashes at the interface with EF-Tu that alter its ability to both engage aminoacylated tRNA termini and undergo the rearrangements needed for stable ternary complex formation. Inspection and in silico analysis of the previously reported structure of EF-Tu(GDPNP)-L-α-Phe-tRNA^Phe^ complex (PDB ID: 1OB2) suggested that all four β-Phe monomers would clash with EF-Tu, while *p*-Az-Phe would not (**Fig. S5**). Reorientation of domains DI-III of EF-Tu, combined with repositioning of the aminoacylated tRNA termini could, in principle, relieve the observed steric clashes with the β-Phe monomers but this change would likely come at the expense of precise positioning of nucleotide binding pocket elements, including the S1 and S2 regions and therefore ternary complex stability (**Fig. 1B**). These findings support the hypothesis that engineering ternary complexes to compensate for nnAA-specific perturbations in stability could significantly improve incorporation efficiencies.

Further in-depth analyses of translation kinetics to address the issue of low non-α-amino acid utilization efficiency is warranted. To date, such studies have been performed under two different experimental approaches: *in vitro* translation^48,75^ or *in cellulo* incorporation^68^. *In vitro* translation systems allow for optimization over a wide range of experimental conditions and have been successful at incorporating diverse non-α-amino acid monomers in short peptides. However, their reaction yields are seldom reported and only a minor fraction of published studies focus on relative rates or mechanistic bottlenecks^16^. *In cellulo* studies that show incorporation of nnAAs outside of α-amino or α-hydroxy acid are far fewer^6–8,76^. Parallel *in vitro* and *in cellulo* investigations are likely to be most informative.

Despite clear evidence that β-Phe monomers cause defects in ternary complex formation and stability, recent experiments show that β^3^-amino or β^2^-hydroxy acids can be incorporated into proteins in cells^6,7,45^. β^3^-(*p*-Br)-Phe has also been incorporated into DHFR in *E. coli* cells expressing a ribosome containing a remodeled PTC^6^. A β^2^-hydroxy analogue of N^ε^-Boc-L-α-Lysine has also been incorporated into sfGFP by a wild-type *E. coli* strain^7^, although the yield of protein was far lower based than expected from the *in vitro* activity of the aaRS. Kinetic simulations of ternary complex formation reveal that this discrepancy is likely explained by the presence of elevated EF-Tu and tRNA concentrations in the cell that help drive ternary complex formation despite significant reductions in stability (**Fig. S4**). Despite this mass action effect, perturbations to the tRNA selection mechanism, and proofreading specifically, are nonetheless expected to remain.

To achieve incorporation efficiencies approximating native amino acids, engineered EF-Tu and tRNA mutants, and potentially other translation machinery components, will likely be required to compensate for monomer-induced penalties to the system. Such efforts should aid the design of tailored orthogonal ternary complex components to complement existing OTS technologies, increasing nnAA tolerance and desired product yields.

## EXPERIMENTAL PROCEDURES

### tRNA^Phe^ purification and fluorescent labeling

Native tRNA^Phe^ was expressed and purified as described previously ^19,36,77^. Briefly, pBS plasmid containing the tRNA^Phe^ V gene was transformed into JM109 cells and incubated overnight at 37 °C with shaking at 250 rpm. Cells pelleted at 10,000 x g for 15 minutes. Cells were lysed by sonication in 20 mM potassium phosphate buffer pH 6.8 with 10 mM Mg(OAc)_2_ and 10 mM β-mercaptoethanol. Cellular debris was pelleted by high-speed centrifugation at 55,000 x g for 1.5 hours. Supernatant was phenol:chloroform extracted twice followed by a series of precipitation steps consisting of an EtOH precipitation, two isopropanol precipitation steps (added dropwise) and a final EtOH precipitation. Bulk tRNA^Phe^ was then aminoacylated for 10 minutes in charging buffer (50 mM Tris-Cl, pH 8.0; 20 mM KCl; 100 mM NH_4_Cl; 10 mM MgCl_2_; 1 mM DTT; 2.5 mM ATP; and 0.5 mM EDTA) with PheRS (crude tRNA in 45-fold molar excess) and a 10-fold molar excess of L-phenylalanine. Phe-tRNA^Phe^ was separated from crude tRNA using a TSK Phenyl-5PW HIC column (Tosoh Bioscience) with a linear gradient starting from buffer A (10 mM NH_4_OAc, pH 5.8; 1.7 M (NH_4_)_2_SO_4_) to buffer B (10 mM NH_4_OAc, pH 5.8; 10% MeOH). Elution fractions corresponding to Phe-tRNA^Phe^ were pooled, dialyzed into storage buffer (10 mM KOAc, pH 6.0; 1 mM MgCl_2_) and concentrated.

Phe-tRNA^Phe^ was fluorescently labeled with Cy^TM^3B Mono NHS ester (Cytiva) using the native aminocarboxypropyluridine (acp^3^) post-transcriptional modification at the U47 position with previously described procedures ^19,36^. Briefly, Phe-tRNA^Phe^ was buffer exchanged into 312.5 mM HEPES, pH 8.0. NaCl was added to a final concentration of 1 M. 2 μL of 50 mM Cy^TM^3B in DMSO was added at 15-minute intervals over a 1-hour period. Reaction conditions were such that all the acyl bond is hydrolyzed during labeling yielding deacyl Cy^TM^3B-tRNA^Phe^. Labeled tRNA^Phe^ was then phenol:chloroform extracted, EtOH precipitated, and purified over the TSK Phenyl-5PW column as described above. Cy3B-tRNA^Phe^ was buffer exchanged into storage buffer, concentrated, aliquoted and stored at −80 °C until further use.

### Elongation factor purification and fluorescent labeling

His-tagged versions of elongation factors Tu and Ts were expressed from pPROEX vectors and purified by Ni^2+^-NTA as described previously ^19,36^. Fluorescent EF-Tu was labeled on a C-terminal acyl-carrier-protein (ACP) tag with LD655-CoA via ACP synthase. Briefly, 5-10 molar equivalents of LD655-CoA was mixed with EF-Tu-ACP in labeling buffer containing 50 mM HEPES pH 7.5, 10 mM Mg(OAc)_2_. Labeled EF-Tu-LD655 was separated from ACP-S and free dye via Ni^2+^-NTA, TEV protease was added to remove the 6X-His tag from EF-Tu-LD655 and run over a second Ni^2+^-NTA to remove TEV. EF-Tu-LD655/Ts complexes were purified by size exclusion chromatography, concentrated into factor storage buffer containing 10 mM HEPES pH 7.5, 100 mM KCl, 1 mM DTT, and 50% glycerol and stored at −20 °C.

### Monomer synthesis

The general procedure for L-α-Phe cyanomethyl ester (CME) monomer synthesis followed previously published procedures with slight modifications ^44^. To a 5-mL round-bottom flask, N-Boc protected amino acid (0.5 mmol) was dissolved in 1 mL of tetrahydrofuran. Flask was then charged with 315 μL of chloroacetonitrile (5.0 mmol, 10 eq.), followed by addition of 100 μL of N,N-diisopropylethylamine (0.6 mmol, 1.2 eq.). The flask was capped with septa and stirred at room temperature overnight, 16 hours. Solvent was removed via rotary evaporation then the crude material was purified by reverse-phase flash chromatography, 0-100% acetonitrile in water, holding at 60% acetonitrile until product was collected. Solvent removed via rotary evaporation, where the resulting oil was dissolved in 1 mL of tetrahydrofuran for deprotection. To the resulting solution, 1.9 mL of trifluoroacetic acid (25 mmol, 50 eq.) was added, and allowed to stir at room temperature for 2 hours. Upon completion, the solvent was removed followed by purification by reverse-phase flash chromatography utilizing a 2% acetonitrile in water mobile phase. Solvent was removed by lyophilization to yield target materials.

The general procedure for β^3^-substituted phenylalanine CME monomers was performed as follows. The Boc-protected-Amino Acid-CME (ca 0.5 mmol) was treated with neat formic acid (2 mL). The solution was stirred at rt for 12 h before removing all the formic acid under reduced pressure by azotropic distillation with CHCl_3_ to afford a pale, yellow oil. The oil was dissolved in minimum amount of THF (ca. 2 mL), triturated with excess MTBE or Et_2_O until white solid was formed persistently. All the residual solvent was removed under reduced pressure. The white solid was crushed into fine powder, rinsed thoroughly with Et_2_O (10 mL) and dried over vacuum for overnight. The typical yield over two steps was 50%.

### Flexizyme charging of native Cy3B-labeled tRNA^Phe^

All concentrations listed are final. 5 mM of CME monomers were charged onto Cy3B-tRNA^Phe^ with a 5-fold excess of flexizyme. Flexizyme RNA oligo sequence which was developed and described previously, ^43,44,78^ used in this study is as follows: GGAUCGAAAGAUUUCCGCGGCCCCGAAAGGGGAUUAGCGUUAGGU. were ordered de-protected from IDT, resuspended in ultra-pure water, flash frozen and stored at −80 C. For L-α-Phe and *p*-Az-Phe, charging reactions were carried out in 100 mM HEPES, pH 6.6, 600 mM MgCl_2_, and 20% DMSO for 2 hours on ice. For the β^2^-substituted Phe monomers, reactions were carried out in 50 mM Bicine, pH 9.0, 600 mM MgCl_2_, and 30% DMSO on ice for 24 hours. For the β^3^-substituted Phe monomers, reactions were carried out in 50 mM Bicine, pH 9.0, 600 mM MgCl_2_, and 10% DMSO on ice for 24 hours. All flexizyme charging reactions were quenched with 90 μL of 0.3 M NaOAc, pH 5.3 and EtOH precipitated at −20 °C. Precipitate was centrifuged at 21,000 x g for 10 minutes, EtOH aspirated, precipitate was resuspended in Buffer A and the acylated species was purified from the deacylated species as described above for normal tRNA^Phe^ purification procedures. Purified, charged monomers were dialyzed into storage buffer and concentrated down via Amicon centrifugal filters with a 3k MWCO. Samples were aliquoted, flash frozen, and stored at −80 °C.

### Ternary complex assay

Ternary complex assays (**Fig. 2** and **3**) were performed with a QuantaMaster-400 Spectrofluorometer (Photon Technology International) with 520 nm and 570 nm excitation and emission wavelengths, as previously described or a micro stopped-flow system (µSFM, BioLogic) equipped with a MOS-200/M spectrometer with the excitation monochromator set at 520 nm to monitor changes in Cy3B fluorescence ^19,36^. In both cases a 532 long bandpass filter was placed in front of the emission PMT to omit noise as a result of excitation light. All concentrations listed are final. Ternary complex formation reactions were carried out in ternary complex reaction buffer containing 100 mM HEPES; pH 7.4, 20 mM KCl, 100 mM NH_4_Cl, 1 mM DTT, 0.5 mM EDTA, and 2.5 mM Mg(OAc)_2_. Briefly, ternary complex formation was achieved by stopped-flow injection of 400 nM (unless specified otherwise) EF-Tu-LD655 into a solution containing 5 nM aa-tRNA^Phe^-Cy3B in ternary complex reaction buffer. Prior to stopped-flow injection, EF-Tu-LD655 was preincubated in ternary complex reaction buffer with 10 µM GTP with or without EF-Ts as indicated. Upon reaction equilibration, ternary complex dissociation was achieved by the stopped-flow injection of 100 μM GDP in ternary complex buffer to the same solution. For the µSFM system, equal volumes of EF-Tu-LD655 with or without 3 µM EF-Ts and aa-tRNA^Phe^-Cy3B were rapidly mixed together (final volume 24 µL, flow rate 1.2 mL/s) the formation reaction was monitored at 800 V with sampling times of 1 ms for the first 5 seconds and 10 ms for the remaining reaction time. All relative fluorescence values were plotted versus time and fit to:

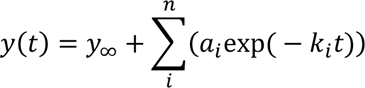

where n = 1 or 2, as required.

### Kinetic simulations

Kinetic analysis of ternary complex formation assays was performed in MATLAB (2021b). *k*_on_ and *k*_off_ rates derived from the µstopped-flow data were used in combination with previously reported values to simulate ternary complex levels at physiological concentrations. A system of ordinary differential equations based on the kinetic model in **Fig. 3C** was solved using the function ode89 in MATLAB at different combinations of aa-tRNA, EF-Tu and EF-Ts concentrations. A proportion of 4*[EF-Tu] = [EF-Ts] was kept constant for each aa-tRNA-EF-Tu pair. Previously reported *E. coli* cytoplasmic concentrations of GDP (0.69 mM) and GTP (4.9 mM) were used. The equilibrium concentrations of aa-tRNA free and bound were used to determine the ternary complex fraction with the following equation:

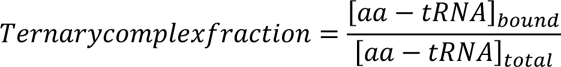

### Single-molecule FRET experiments

Single-molecule FRET experiments were performed using a custom-built, prism-type TIRF microscope. Bacterial ribosomes programed with Cy3-fMet-tRNA^fMet^ in the P site and a UUC codon displayed in the A site were surface immobilized via streptavidin-biotin interaction to a transparent surface passivated with polyethylene glycol (PEG) polymers doped with biotin-PEG. tRNA selection experiments were initiated by injection of pre-formed ternary complexes (aa-tRNA-LD655, 12.5 nM; EF-Tu/Ts 125 nM; GTP or GTPγS 500 µM) in bacterial polymix buffer (50 mM Tris-OAc; pH 7.5, 100 mM KCl, 5 mM NH_4_OAc, 0.5 mM Ca(OAc)_2_, 5 mM Mg(OAc)_2_, 6 mM 2-mercaptoethanol, 0.1 mM EDTA, 5 mM putrescine and 1 mM spermidine) supplemented with 2 mM PCA/PCD oxygen scavenging system and 1 mM each of cyclooctatetraene (COT), nitrobenzyl alcohol (NBA), and Trolox triplet state quenchers. Translocation experiments were initiated by injection of pre-formed ternary complexes (aa-tRNA-LD655, 25 nM; EF-Tu/Ts 250 nM; GTP 1250 µM) supplemented with 10 µM EF-G in bacterial polymix buffer. Samples were illuminated with a 532 nm diode pumped solid-state laser (Opus, LaserQuantum) at 2.0 and 0.25 kW cm^-2^ (0.16 kW cm^-2^ for translocation experiments) with 10 or 100-ms integration times, respectively. Fluorescence emission from donor and acceptor fluorophores was collected using a 60X/1.27 NA super-resolution water-immersion objective (Nikon). Fluorescence was recorded onto two aligned ORCA-Fusion sCMOS cameras (C-14440-20UP, Hamamatsu). Instrument control was performed using custom software written in LabVIEW (National Instruments). Fluorescence intensities were extracted from the recorded videos and FRET efficiency traces were calculated using the SPARTAN software package. FRET traces were selected for further analysis according to the following criteria: 8:1 signal/background-noise ration and 6:1 signal/signal-noise ratio, less than four donor-fluorophore blinking events and a correlation coefficient between donor and acceptor of <0.5. The resulting smFRET traces were further post synchronized to the appearance of FRET and analyzed using the segmental *k-*means (SKM) algorithm as implemented in the SPARTAN software package v3.8. Data were plotted in OriginPro 2019b (OriginLab, Northhampton, MA). Dwell time curves were fit to:

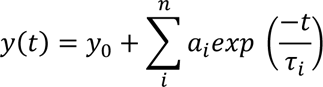

where n = 1 or 3 as required. Mean and standard deviations of classical-hybrid split ratio were calculated from four independent measurements. Statistical significance (*p* < 0.05) was assessed by a two-way ANOVA followed by a *post-hoc* Bonferroni test.

## AUTHOR CONTRIBUTIONS

A.S., S.J.M. and S.C.B. conceived the project. F.A.C-N. and W.C.G. expressed, purified and labelled tRNA and EF-Tu, performed and analyzed ensemble and single-molecule FRET measurements on tRNA selection. W.C.G. carried out flexizyme charging of dye-labelled tRNA. Y-C.C. and I.K. synthetized CME-activated L-α-Phe and β-Phe monomers. M.M. synthetized the *p*-Az-Phe monomer. J.A. performed single-molecule FRET measurements on translocation and assisted with data analysis. S.K.N. performed molecular modelling. R.B.A. assisted with the synthesis of the fluorophores employed. F.A.C-N., W.C.G, J.A., A.S. and S.C.B. wrote the manuscript with input from all the authors.

## ACKNOWLEDGMENTS

This work was supported primarily by the National Science Foundation Center for Genetically Encoded Materials (C-GEM), an NSF Center for Chemical Innovation (NSF CHE-2002182; F.A.C-N., W.C.G., Y-C.C., I.K., A.S., S.J.M., and S.C.B.) C-GEM funding supported protein and tRNA expression, reagent synthesis and purification, chemistry and biochemistry efforts associated with all assays reported, ensemble and single-molecule fluorescence investigations, and manuscript preparation. Additional support was provided by the National Institutes of Health (5R01AI150560; M.M.), primarily for the synthesis of the *p*-Az-Phe monomer and the fluorophores employed. We thank St. Jude Children’s Research Hospital for their support of J.A., S.K.N. and R.B.A and the Single-Molecule imaging Center, Daniel S. Terry, Zeliha Kilic and Mikael Holm in particular, for early guidance with kinetic simulations, their training and their thoughtful discussions and comments during manuscript preparation.

## CORRESPONDING AUTHORS

Scott C. Blanchard – Department of Structural Biology, St Jude Children’s Research Hospital, Memphis, Tennessee 38105, United States; Department of Chemical Biology & Therapeutics, St Jude Children’s Research Hospital, Memphis, Tennessee 38105, United States.

Scott J. Miller – Department of Chemistry, Yale University, New Haven, Connecticut 06520-8107 United States.

## SUPPLEMENTAL METHODS

**Synthesis of Boc-Protected β**^2^**-amino acid (Both enantiomeric forms)**

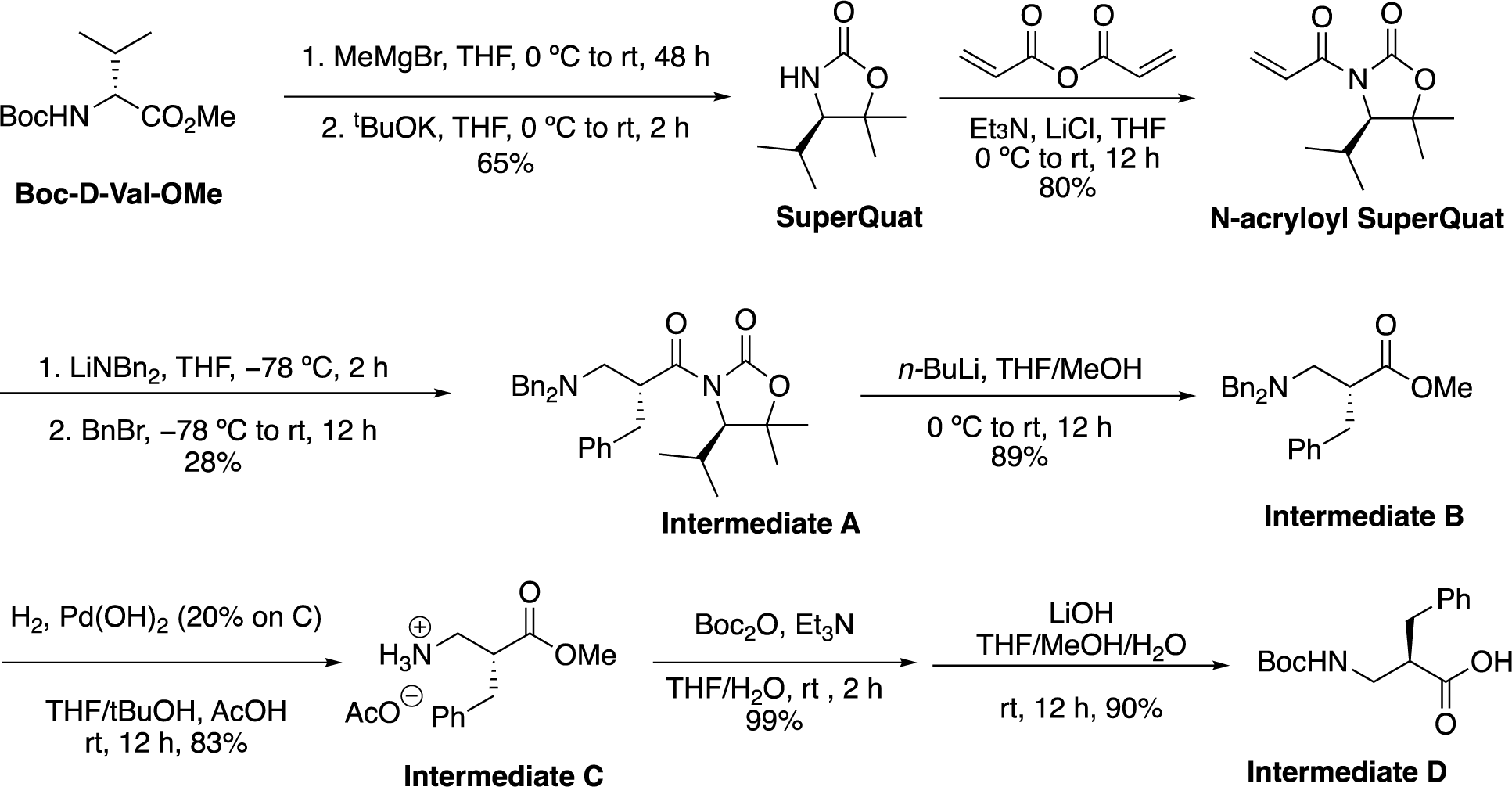

Note: The enantiomer of **Intermediate D** was prepared from Boc-L-Val-OMe.

**SuperQuat** and **N-acryloyl SuperQuat** were prepared according to literature procedure (Asymmetric synthesis of b2-amino acids: 2-substituted-3-aminopropanoic acids from N-acryloyl SuperQuat derivatives. *Org. Biomol. Chem*., **2007**, *5*, 2812–2825.) **Intermediate A** and **B** were also prepared according to the same literature. For **Intermediate A**, we used a different eluent system (hexanes:EtOAc 10: 1 to 7:1, instead of hexanes:Et_2_O 20:1 in the literature condition) for the SiO_2_ column chromatographic purification to afford **Intermediate A** as pale yellow oil (980 mg, 1.96 mmol) in 28% yield (from 7 mmol **N-acryloyl SuperQuat**). For **Intermediate B**, same literature procedure was followed to afford **Intermediate B** as colorless oil (650 mg,1.74 mmol) in 89% yield (from 1.96 mmol of **A**).

Note: The enantiomer of **Intermediate A** and **B** were prepared from Boc-L-Val-OMe as the starting material by the same procedure.

**Intermediate A**

**(*R*)-3-((*R*)-2-benzyl-3-(dibenzylamino)propanoyl)-4-isopropyl-5,5-dimethyloxazolidin-2-one**

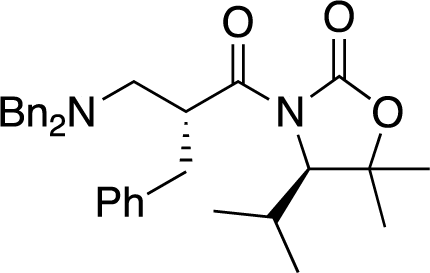

^1^H NMR (400 MHz, CD_2_Cl_2_) δ 7.42 – 7.26 (m, 15H), 4.94 (tt, *J* = 8.6, 6.0 Hz, 1H), 4.23 (d, *J* = 2.8 Hz, 1H), 3.73 (d, *J* = 13.6 Hz, 2H), 3.65 (d, *J* = 13.7 Hz, 2H), 3.11 – 2.89 (m, 3H), 2.59 (dd, *J* = 12.6, 5.6 Hz, 1H), 2.05 (pd, *J* = 6.9, 2.8 Hz, 1H), 1.53 (d, *J* = 3.0 Hz, 6H), 0.83 (d, *J* = 7.0 Hz, 3H), 0.69 (d, *J* = 6.8 Hz, 3H).

^13^C NMR (101 MHz, CD_2_Cl_2_) δ 175.6, 153.8, 139.6, 139.4, 129.7, 129.6, 128.8, 128.5, 127.3, 126.7, 82.7, 66.7, 58.9, 56.6, 43.1, 37.6, 29.9, 29.1, 21.5, 21.4, 16.6.

***enantiomer*-Intermediate A**

**(*S*)-3-((*S*)-2-benzyl-3-(dibenzylamino)propanoyl)-4-isopropyl-5,5-dimethyloxazolidin-2-one**

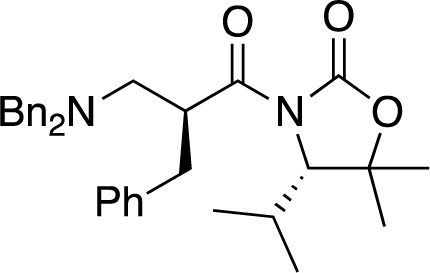

^1^H NMR (500 MHz, CD_2_Cl_2_) δ 7.39 – 7.18 (m, 15H), 4.84 (tt, *J* = 8.5, 6.0 Hz, 1H), 4.16 (d, *J* = 2.8 Hz, 1H), 3.65 (d, *J* = 13.6 Hz, 2H), 3.58 (d, *J* = 13.6 Hz, 2H), 3.02 – 2.90 (m, 2H), 2.85 (dd, *J* = 13.6, 8.8 Hz, 1H), 2.51 (dd, *J* = 12.6, 5.7 Hz, 1H), 2.06 – 1.95 (m, 1H), 1.49 (d, *J* = 13.8 Hz, 6H), 0.77 (d, *J* = 7.0 Hz, 3H), 0.63 (d, *J* = 6.8 Hz, 3H).

^13^C NMR (126 MHz, CD_2_Cl_2_) δ 175.6, 153.8, 139.6, 139.5, 129.6, 129.5, 128.7, 128.5, 127.3, 126.6, 82.8, 66.8, 58.9, 56.5, 43.1, 37.6, 29.9, 29.1, 21.5, 21.3, 16.6.

For **Intermediate B** to **Intermediate C**, we modified the condition (Literature: *Org. Biomol. Chem*., **2007**, *5*, 2812–2825.) from Pd/C in MeOH/AcOH into Pd(OH)_2_/C (10% by weight w.r.t. substrate) in THF/*t-*BuOH (v:v, 3:1, 0.1 M) with AcOH (10% by volume) as additive.

**Intermediate B** (650 mg, 1.74 mmol) was dissolved in THF/*t-*BuOH (v:v, 3:1, total 16 mL, *ca*. 0.1 M) and AcOH (1.6 mL). Pd(OH)_2_ (20% on carbon) (65 mg) was added to the solution. The reaction was degassed, connected to a H_2_ balloon and stirred at rt for overnight before filtration over Celite. The filtrate was concentrated under reduced pressure to afford **Intermediate C** as oil (364 mg, 1.44 mmol, 83%) which was subjected to next step without further purification. Same experimental sequence was carried out to prepare the enantiomer of **Intermediate C**.

For **Intermediate C** to **Intermediate D**, **C** (364 mg, *ca*. 1.44 mmol, 1 equiv.) was dissolved in THF/H_2_O (6 mL:2 mL). Et_3_N (0.6 mL, 4.3 mmol, 3 equiv.) and Boc_2_O (377 mg, 1.7 mmol, 1.2 equiv.) were added. The solution was stirred at rt for 2 h before dilution with EtOAc (10 mL). The organic phase was washed successively with 10% citric acid solution, saturated NaHCO_3_ solution and brine. The organic phase was dried over anhydrous Na_2_SO_4_, filtered and concentrated to afford (*R*)-Boc-β2-Phe-OMe (423 mg, 1.44 mmol), which was subjected to next step without further purification. LiOH•H_2_O (242 mg, 7.2 mmol, 4 equiv.) and THF/MeOH/H_2_O (4 mL/1 mL/1 mL) were added to the crude product. The mixture was stirred at rt for overnight before dilution with 4 mL H_2_O. The aqueous phase was washed against Et_2_O (5 mL × 2), followed by acidifying with 1N HCl to adjust the pH into ∼2. The aqueous phase was extracted with EtOAc (15 mL × 2), dried over anhydrous Na_2_SO_4_, filtered and concentrated under reduced pressure to afford **Intermediate D** (363 mg, 1.3 mmol, 90%) as a colorless oil. The same experimental sequence was carried out to prepare the enantiomer of **Intermediate D**.

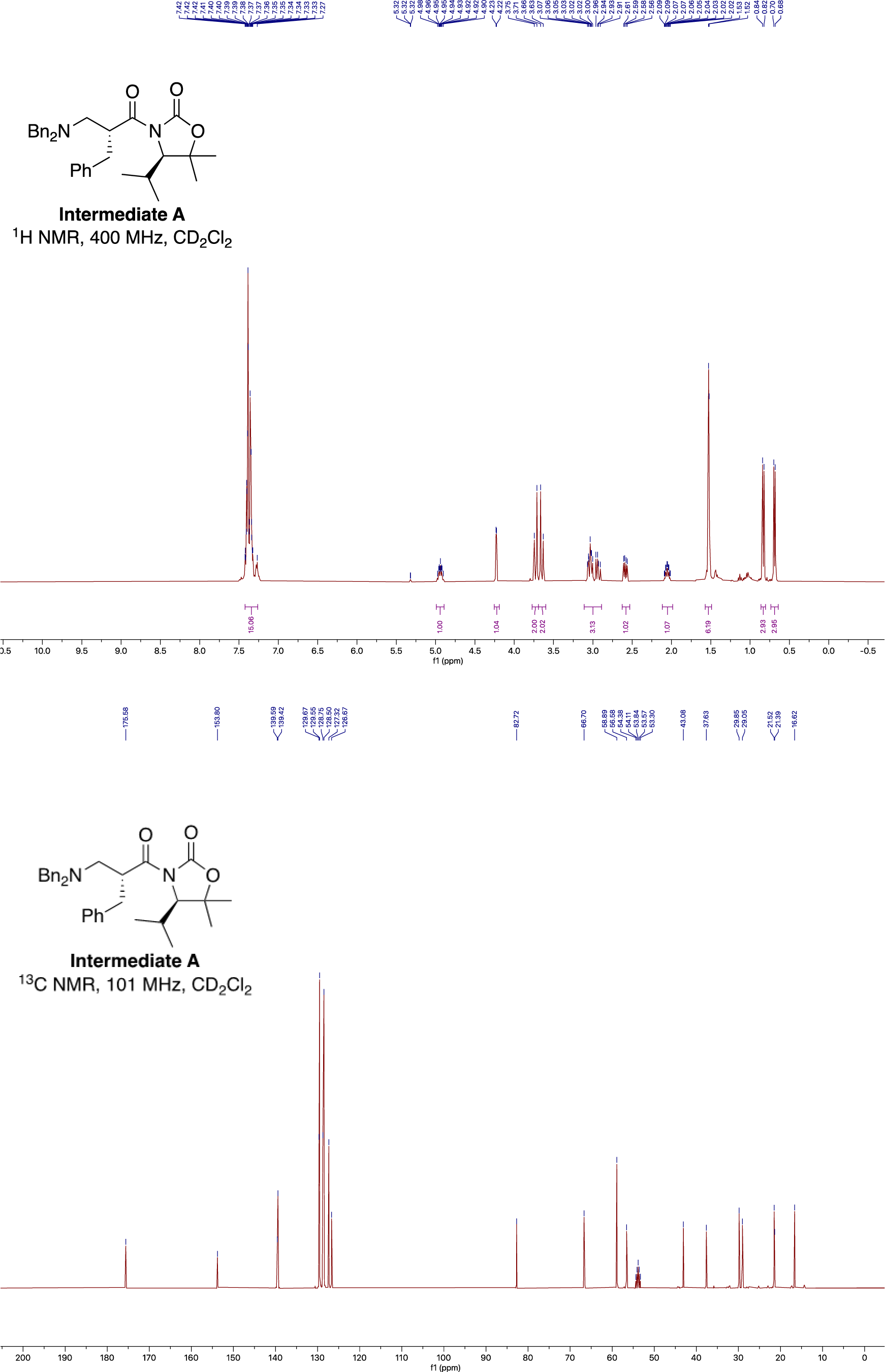
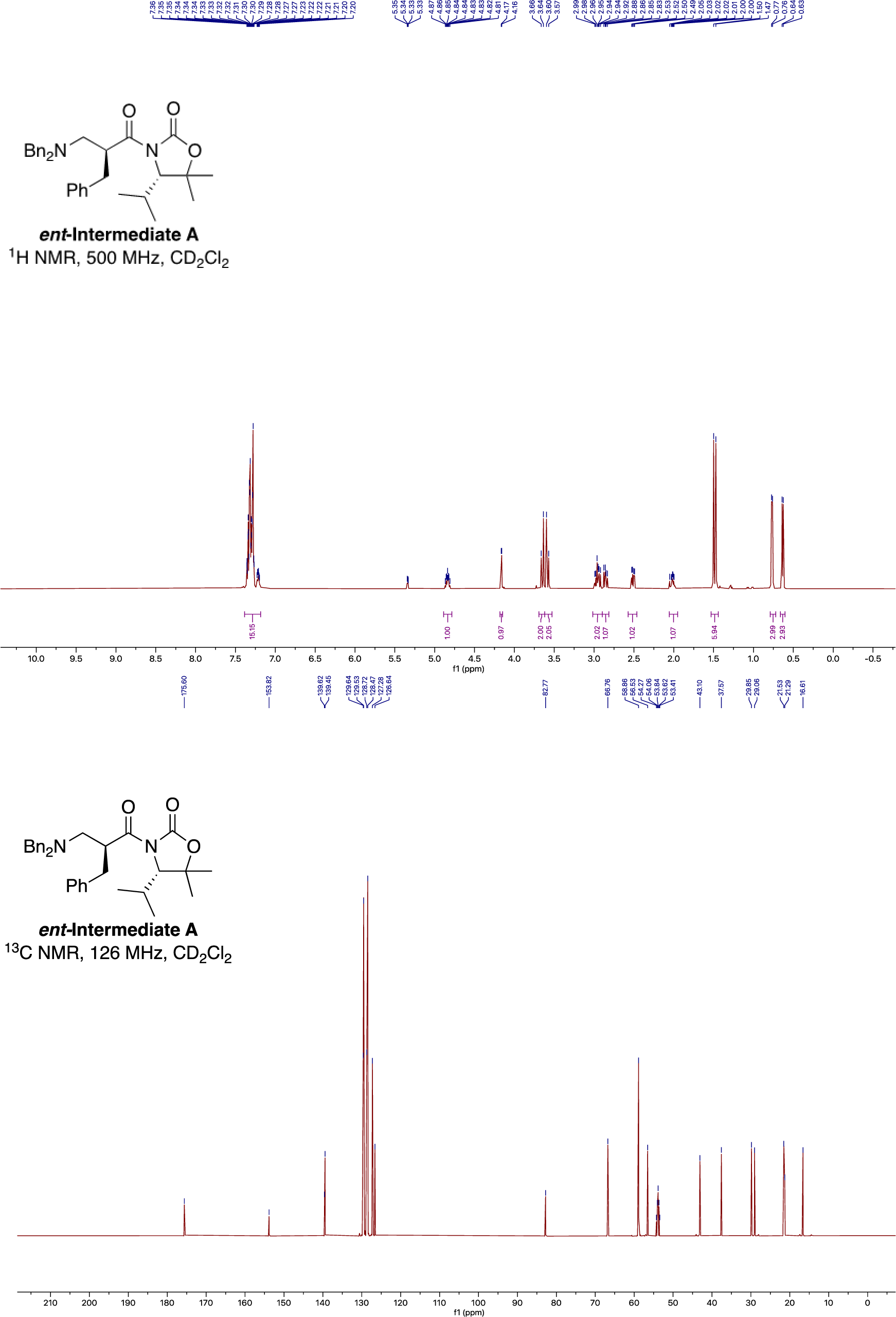

**General procedure for synthesis of β-amino acid-cyanomethyl ester-formate salt**

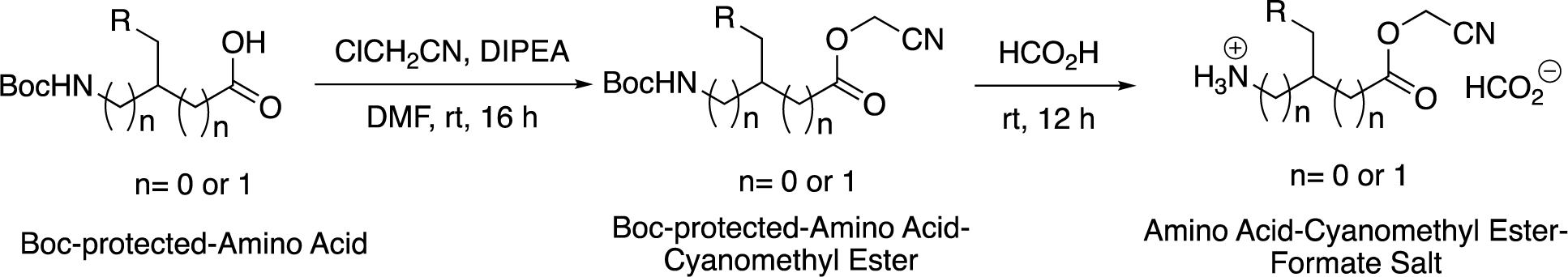

To a stirred solution of Boc-protected-amino acid (0.5 mmol, 1 equiv.) was added dry DMF (1 mL), ClCH_2_CN (0.75 mmol, 1.5 equiv.) and dry DIPEA (1 mmol, 2 equiv.), the solution was stirred at rt for 16 h before dilution with EtOAc (10 mL). The organic phase was washed successively with 10% citric acid aq. solution, 5 % LiCl aq. solution and brine. The organic phase was dried over anhydrous Na_2_SO_4_, filtered and concentrated under reduced pressure to afford the corresponding Boc-protected-Amino Acid-Cyanomethyl ester, which was used without further purification. Note: The cyanomethyl ester is very sensitive to aqueous basic medium.

The Boc-protected-Amino Acid-Cyanomethyl ester (*ca* 0.5 mmol) was treated with neat formic acid (2 mL). The solution was stirred at rt for 12 h before removing all the formic acid under reduced pressure by azeotropic distillation with CHCl_3_ to afford a pale-yellow oil. The oil was dissolved in minimum amount of THF (*ca.* 2 mL), triturated with excess MTBE or Et_2_O until white solid was formed persistently. All the residual solvent was removed under reduced pressure. The white solid was crushed into fine powder, rinsed thoroughly with Et_2_O (10 mL) and dried over vacuum for overnight. The typical yield over two steps was 50%. Note: The final product is very sensitive to water and alcohol which cause saponification or transesterification.

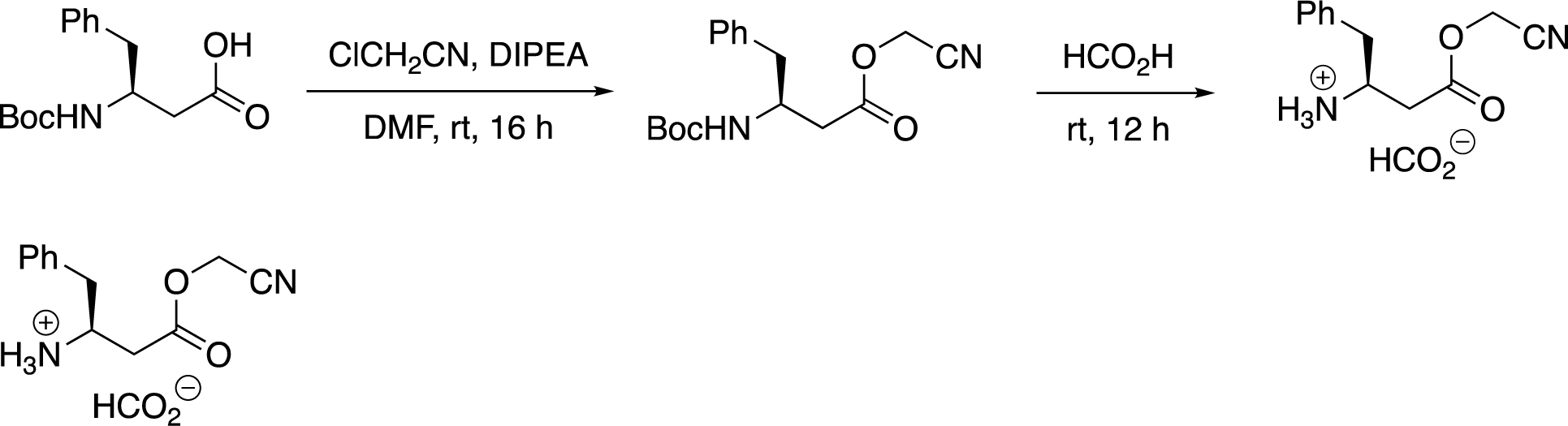

**(*S*)-4-(cyanomethoxy)-4-oxo-1-phenylbutan-2-aminium formate**

^1^H NMR (500 MHz, CDCl_3_) δ 9.22 (s, 3H), 8.37 (s, 1H), 7.41 – 7.26 (m, 5H), 4.72 (s, 2H), 3.89 (s, 1H), 3.23 (dd, *J* = 14.0, 5.6 Hz, 1H), 3.04 – 2.74 (m, 3H).

^13^C NMR (126 MHz, CDCl_3_) δ 169.9, 168.2, 135.4, 129.5, 129.2, 127.7, 114.4, 49.4, 49.1, 39.3, 35.8.

HRMS-ESI: calculated for [C_12_H_15_N_2_O_2_]^+^ 219.1128, found 219.1150.

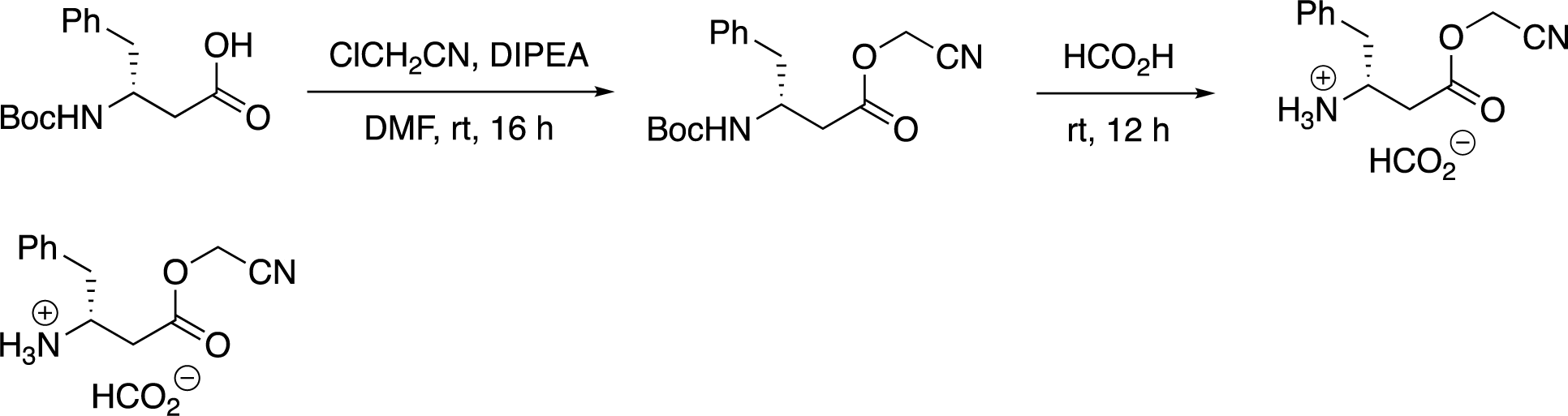

**(*R*)-4-(cyanomethoxy)-4-oxo-1-phenylbutan-2-aminium formate**

^1^H NMR (500 MHz, CDCl_3_) δ 9.15 (s, 3H), 8.36 (s, 1H), 7.43 – 7.20 (m, 5H), 4.70 (s, 2H), 3.87 (s, 1H), 3.22 (dd, *J* = 13.6, 5.6 Hz, 1H), 3.01 – 2.70 (m, 3H).

^13^C NMR (126 MHz, CDCl_3_) δ 169.8, 168.3, 135.5, 129.5, 129.2, 127.7, 114.4, 49.3, 49.0, 39.3, 35.9.

HRMS-ESI: calculated for [C_12_H_15_N_2_O_2_]^+^ 219.1128, found 219.1144.

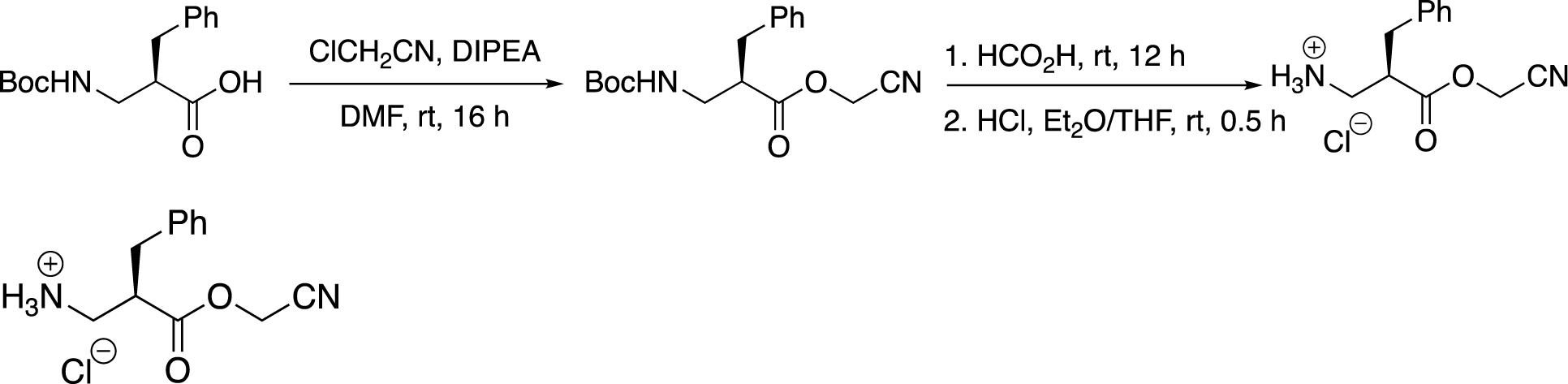

**(*R*)-2-benzyl-3-(cyanomethoxy)-3-oxopropan-1-aminium chloride**

Procedure for anion metathesis with chloride

After removal of residual formic acid from the Boc deprotection, the oily form product (formate salt) was dissolved in minimum amount of THF (*ca* 2 mL). HCl (1M in Et_2_O) (1.5 mL, 3 equiv.) was added. The solution was stirred for 0.5 h before concentration to dryness, and repetitively triturated with excess MTBE or Et_2_O until white solid was formed persistently. All the residual solvent was removed under reduced pressure. The white solid was crushed into fine powder, rinsed thoroughly with Et_2_O (10 mL) and dried over vacuum for overnight.

^1^H NMR (500 MHz, DMSO-*d*6) δ 8.45 (s, 3H), 7.35 – 7.14 (m, 5H), 4.95 (s, 2H), 3.21 (ddq, *J* = 8.3, 6.0, 3.0 Hz, 1H), 3.08 – 2.96 (m, 2H), 2.90 (dd, *J* = 13.8, 8.0 Hz, 2H).

^13^C NMR (126 MHz, DMSO-*d*6) δ 171.1, 137.3, 128.9, 128.5, 126.8, 115.6, 49.6, 44.3, 39.1. HRMS-ESI: calculated for [C_12_H_15_N_2_O_2_]^+^ 219.1128, found 219.1141.

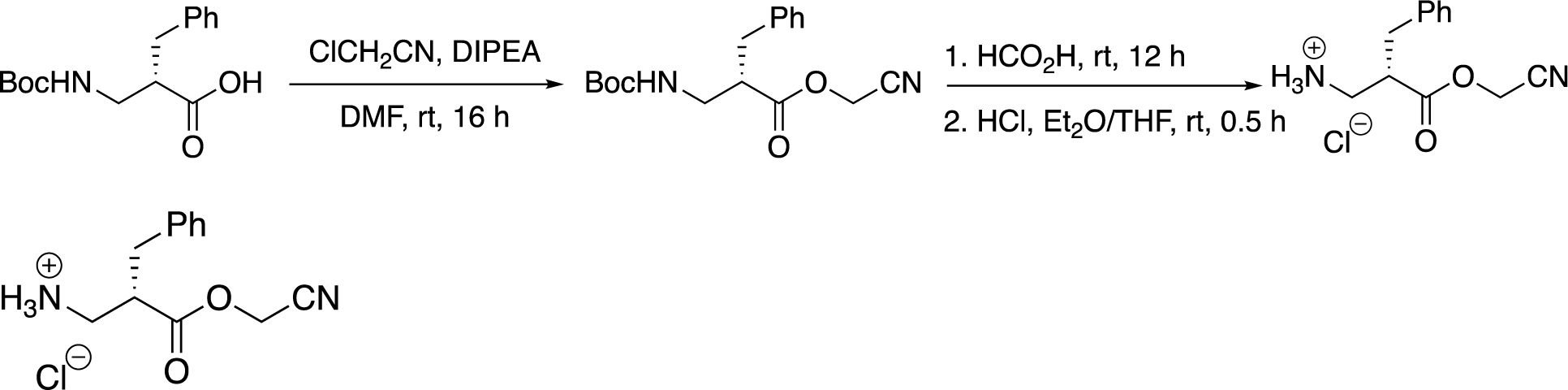

**(*S*)-2-benzyl-3-(cyanomethoxy)-3-oxopropan-1-aminium chloride**

Following the procedure for anion metathesis with chloride.

^1^H NMR (500 MHz, DMSO-*d*6) δ 8.37 (s, 3H), 7.32 – 7.20 (m, 5H), 4.95 (s, 2H), 3.19 (td, *J* = 8.2, 4.1 Hz, 1H), 3.06 – 2.87 (m, 4H).

^13^C NMR (126 MHz, DMSO-*d*6) δ 171.1, 137.3, 128.9, 128.5, 126.8, 115.7, 49.6, 44.4, 39.1. HRMS-ESI: calculated for [C_12_H_15_N_2_O_2_]^+^ 219.1128, found 219.1137.

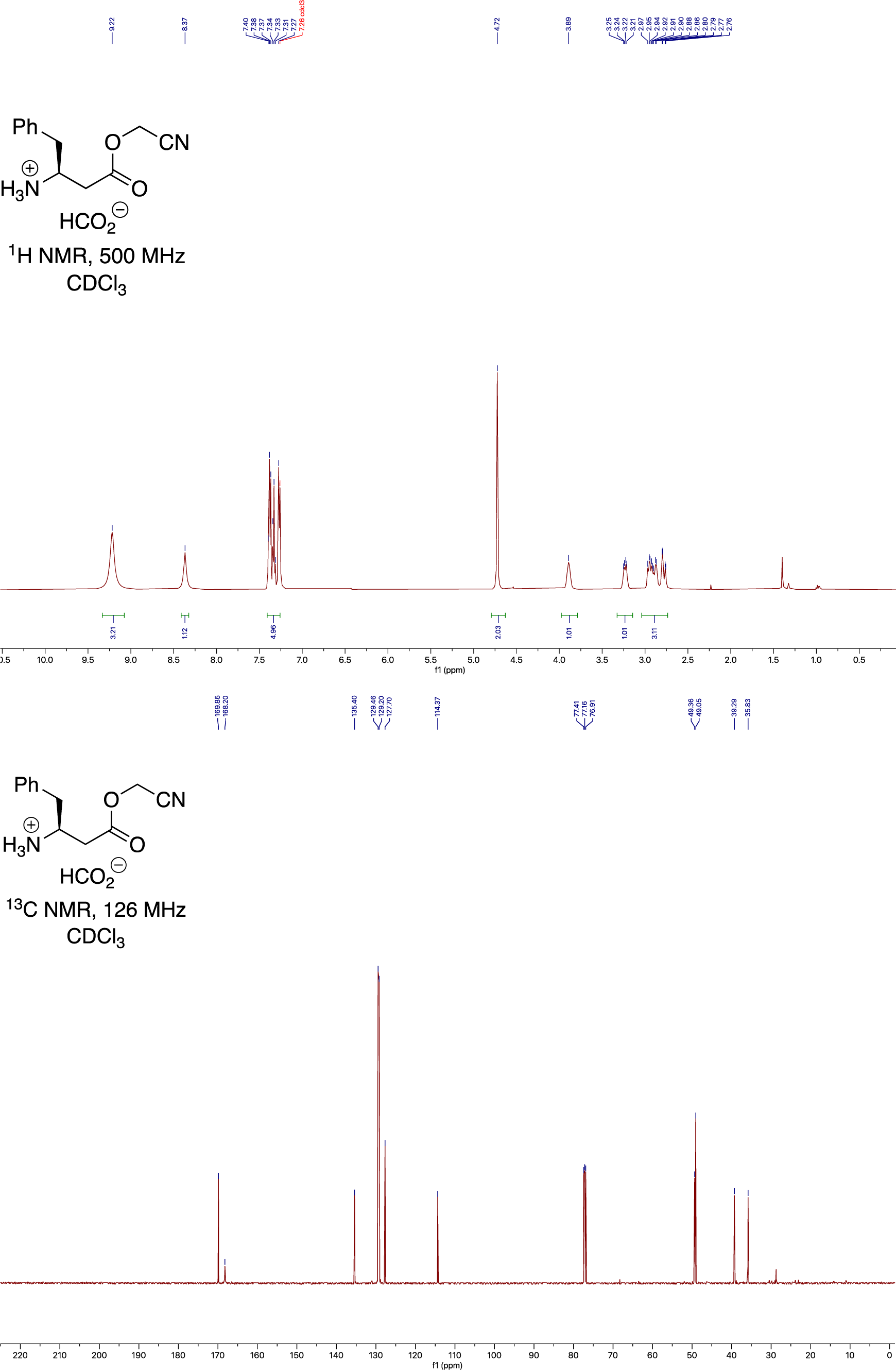
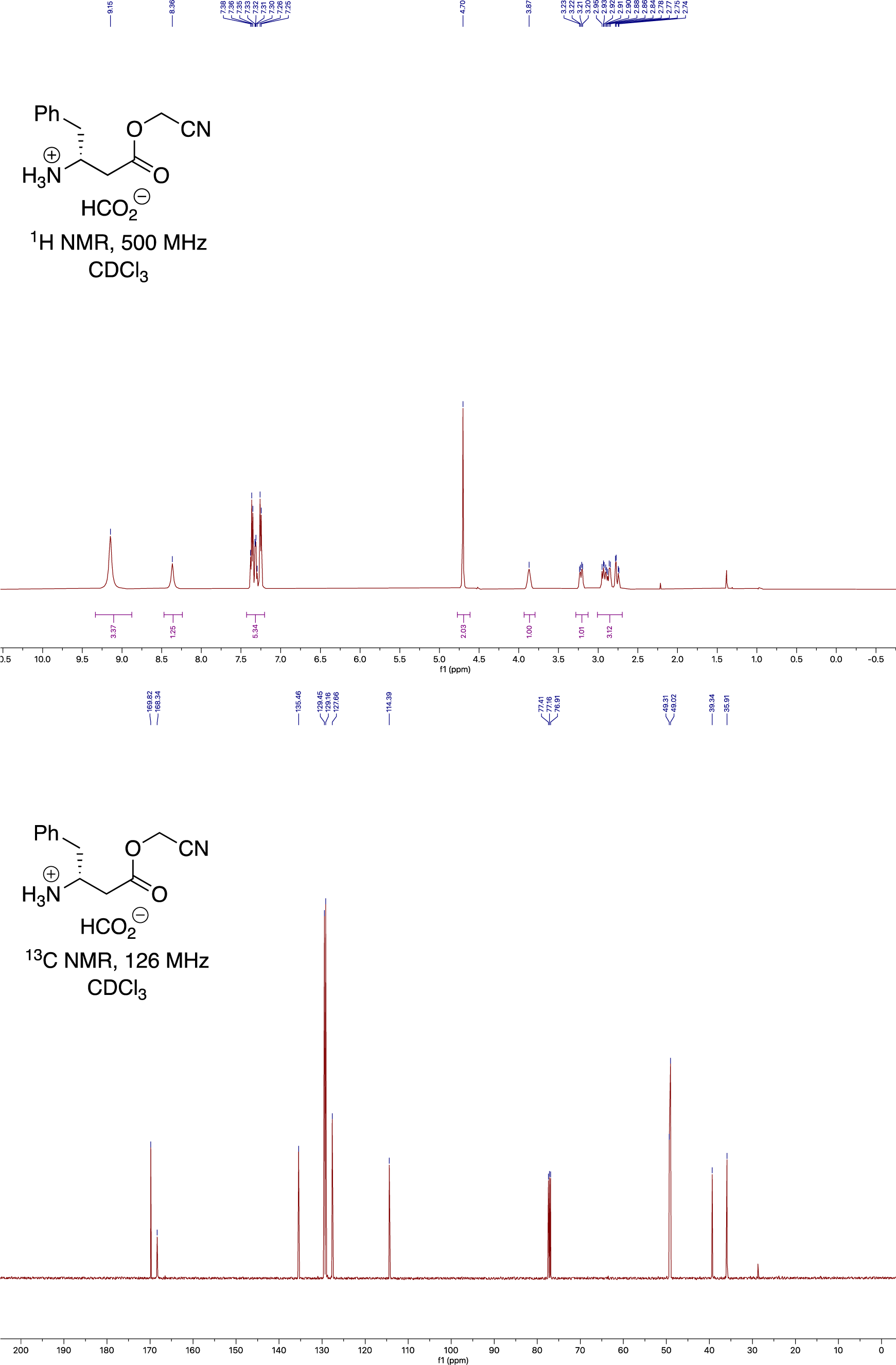
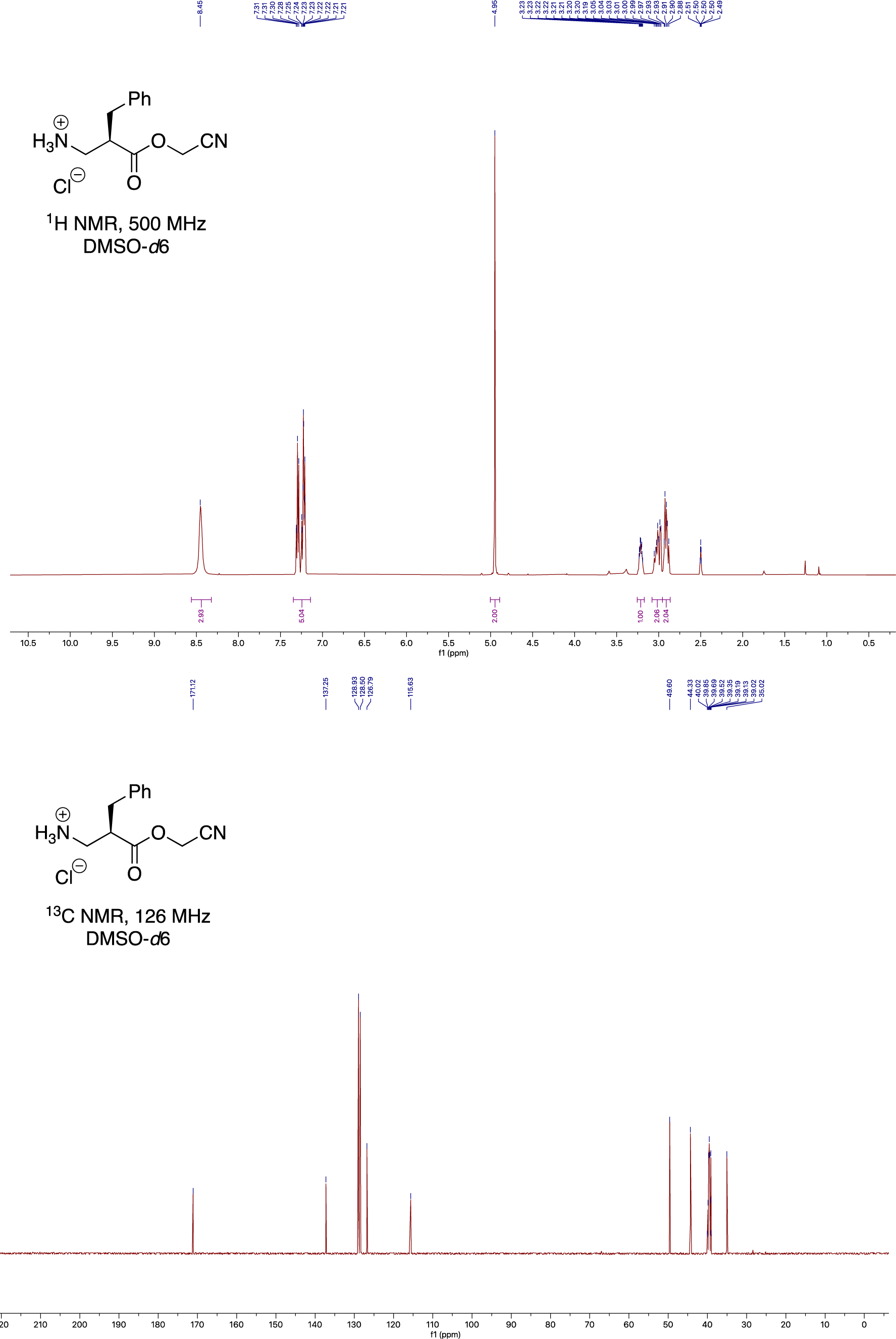
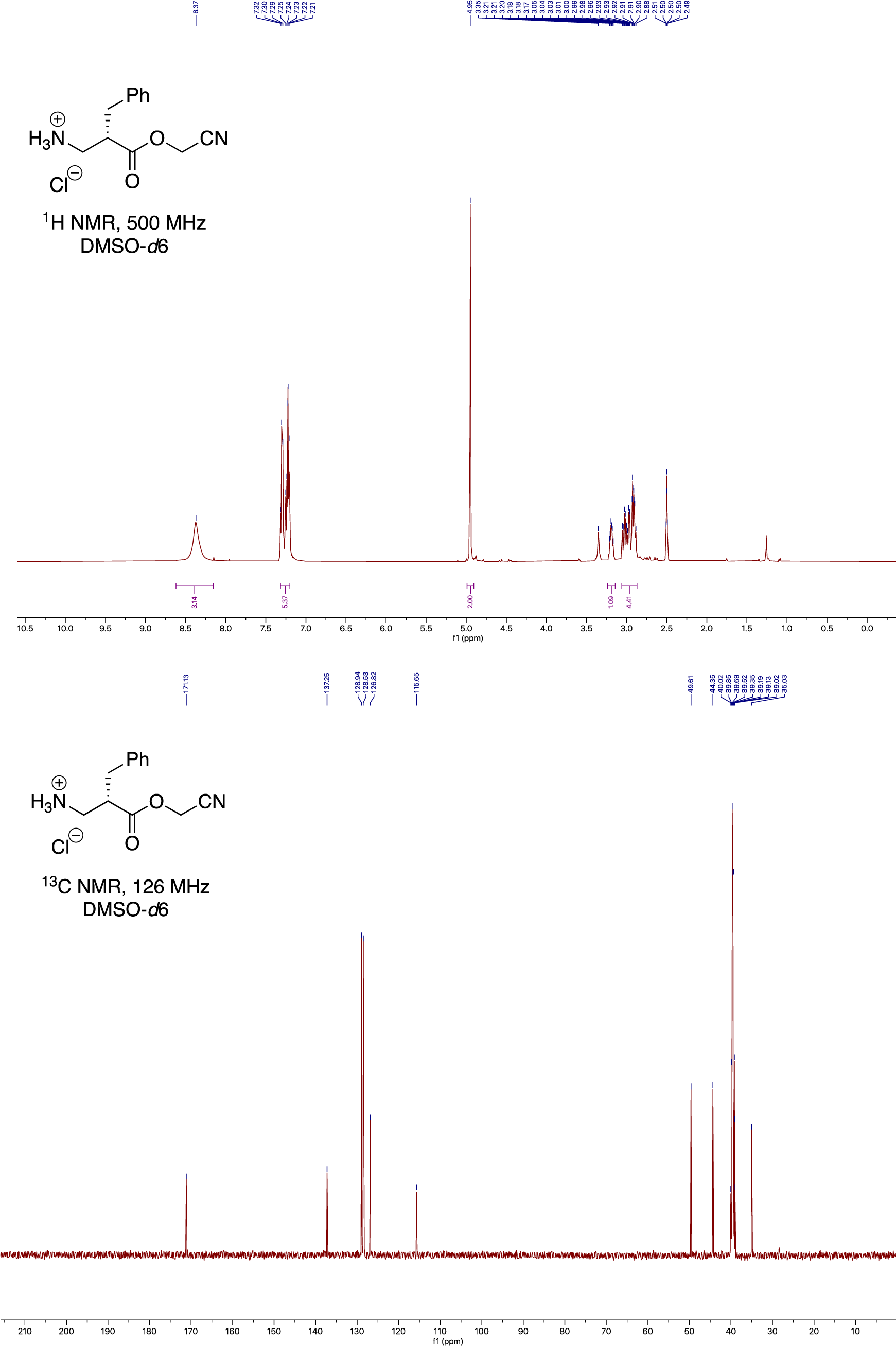

**General procedure for synthesis of α-amino acid-cyanomethyl esters**

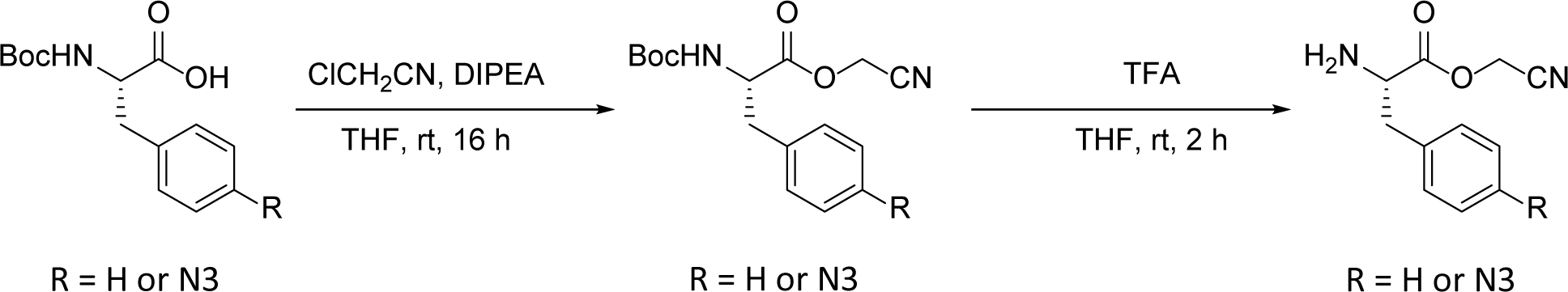

To a 5-mL round-bottom flask, N-Boc protected amino acid (0.5 mmol) was dissolved in 1 mL of tetrahydrofuran. Flask was then charged with 315 μL of chloroacetonitrile (5.0 mmol, 10 eq.), followed by addition of 100 L of N,N-diisopropylethylamine (0.6 mmol, 1.2 eq.). Flask was capped with septa and stirred at room temperature overnight, 16 hours. Solvent was removed via rotary evaporation then the crude material was purified by reverse-phase flash chromatography, 0-100% acetonitrile in water, holding at 60% acetonitrile until product was collected. Solvent removed via rotary evaporation, where the resulting oil was dissolved in 1 mL of tetrahydrofuran for deprotection. To the resulting solution, 1.9 mL of trifluoroacetic acid (25 mmol, 50 eq.) was added, and allowed to stir at room temperature for 2 hours. Upon completion, the solvent was removed followed by purification by reverse-phase flash chromatography utilizing a 2% acetonitrile in water mobile phase. Solvent was removed by lyophilization to yield target materials as colorless oils at 54% and 49% yield for phenylalanine and 4-azidophenylalanine derivatives respectively. Note: The final product is very sensitive to water and alcohol which cause saponification or transesterification.

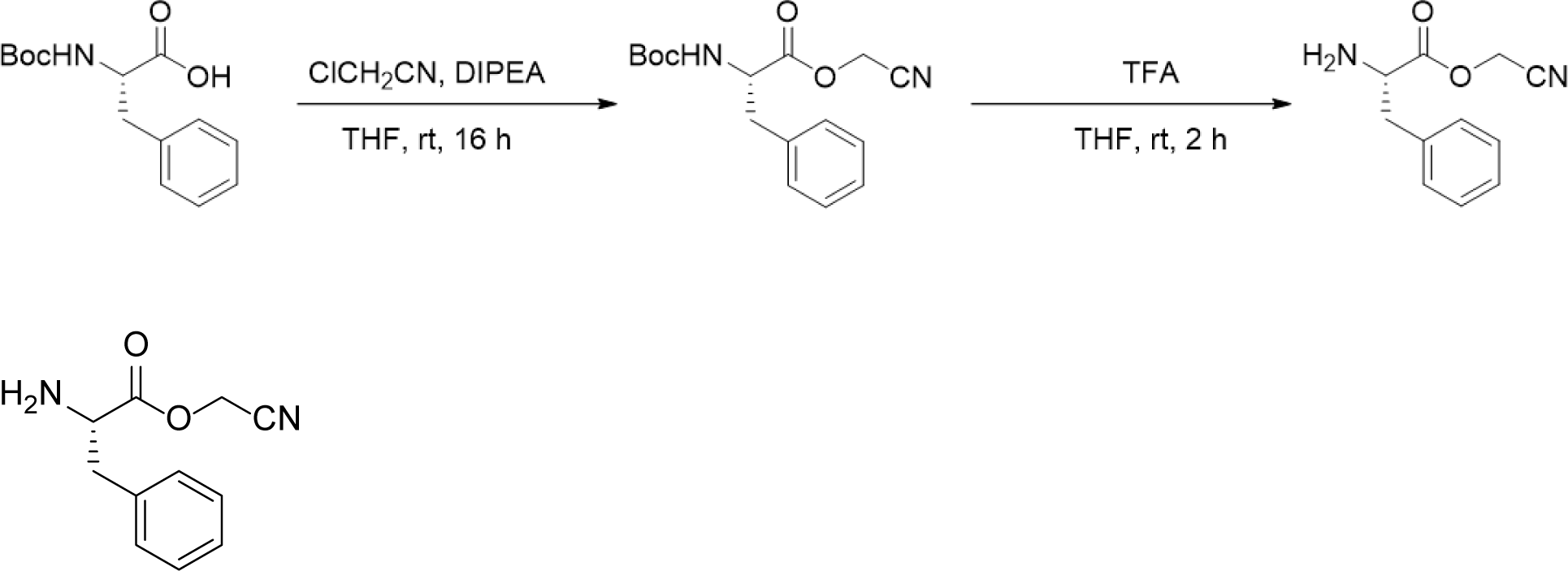

**Cyanomethyl-ester-L-phenylalanine**

^1^H NMR (400 MHz, DMSO-*d*6) δ 8.58 (s, 2H), 7.33 (m, 5H), 5.10 (d, 2H), 4.47 (t, 1H), 3.12 (m, 2H). ESI-MS: calculated for [C_11_H_13_N_2_O_2_]^+^ = 205.10, found 205.24.

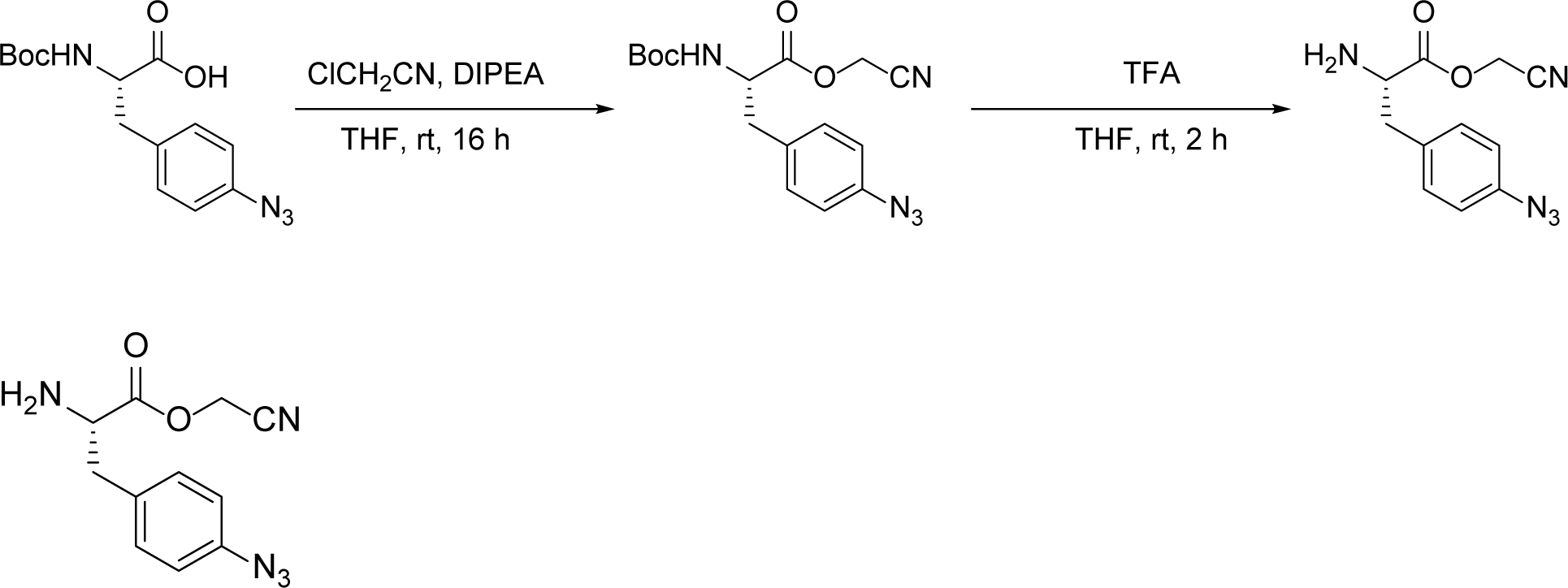

**4-Azido-Cyanomethyl-Ester-L-Phenylalanine**

^1^H NMR (400 MHz, DMSO-*d*6) δ 8.45 (s, 2H), 7.29 (d, 2H), 7.09 (d, 2H), 5.09 (d, 2H), 4.43 (t, 1H), 3.11 (m, 2H). ESI-MS: calculated for [C_11_H_12_N_5_O_2_]^+^ = 246.10, found 246.15.

## SUPPLEMENTAL FIGURE LEGENDS

**Supplemental Figure S1.**
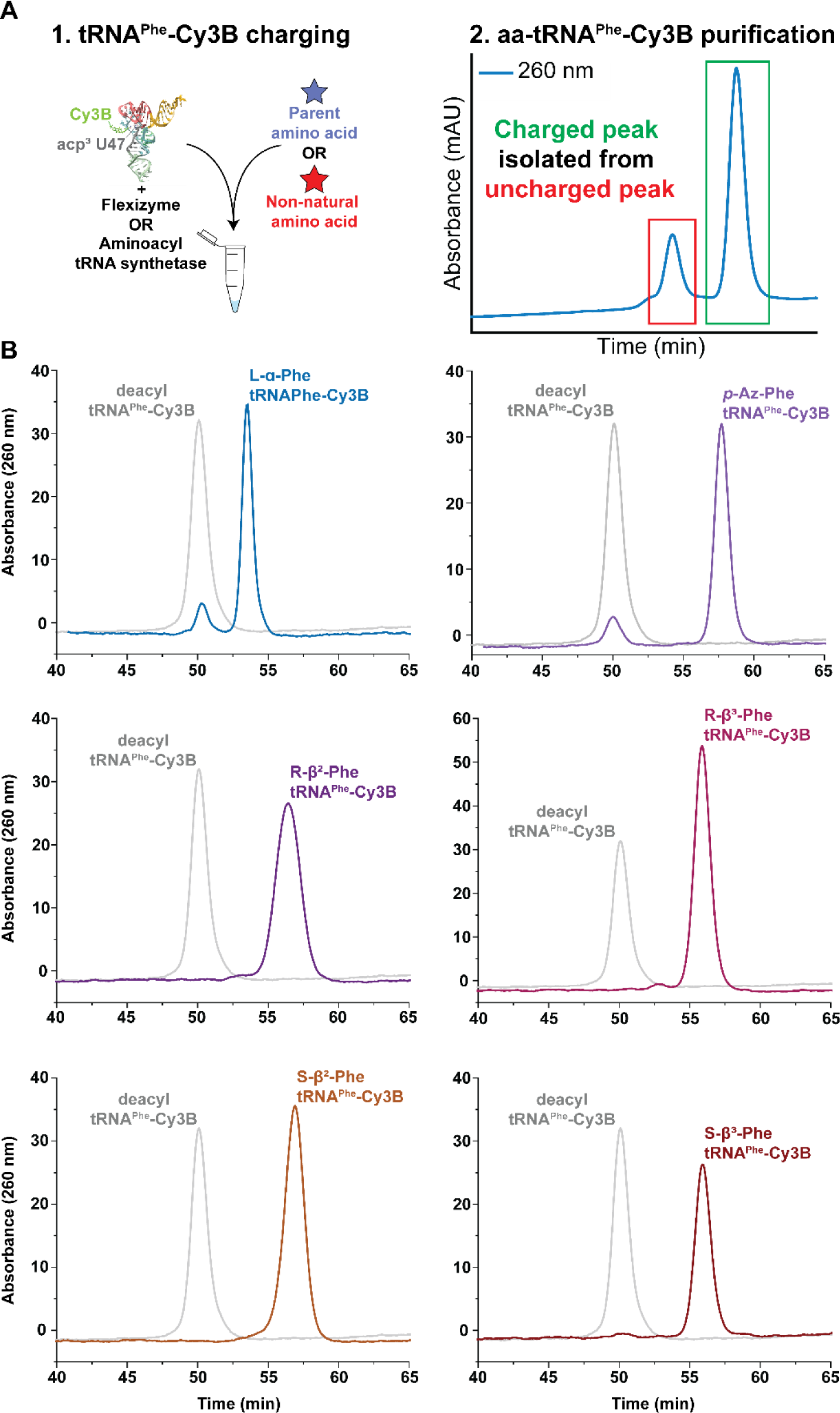
Aminoacylation analysis of various aa-tRNA^Phe^-Cy3B. **A)** Cartoon schematic of tRNA^Phe^-Cy3B charging and purification workflow (see methods for details). De-acylated tRNA^Phe^-Cy3B (variegated) is aminoacylated **(1)** mixed with an excess of flexizyme or tRNA synthetase. Addition of the specified parent (blue) or non-natural (red) amino acid initiates the acylation reaction and is allowed to proceed for the indicated time detailed in the Methods section. Aminoacylated species are then prepped and purified by hydrophobic interaction (HIC) **(2)** by monitoring the absorbance at 260 nm as outlined in the Methods. **B)** Representative HIC purification chromatograms of various parent and non-natural-tRNA^Phe^-Cy3B (various colors and indicated monomers) compared to deacyl-tRNA^Phe^-Cy3B (grey).

**Supplemental Figure S2.**
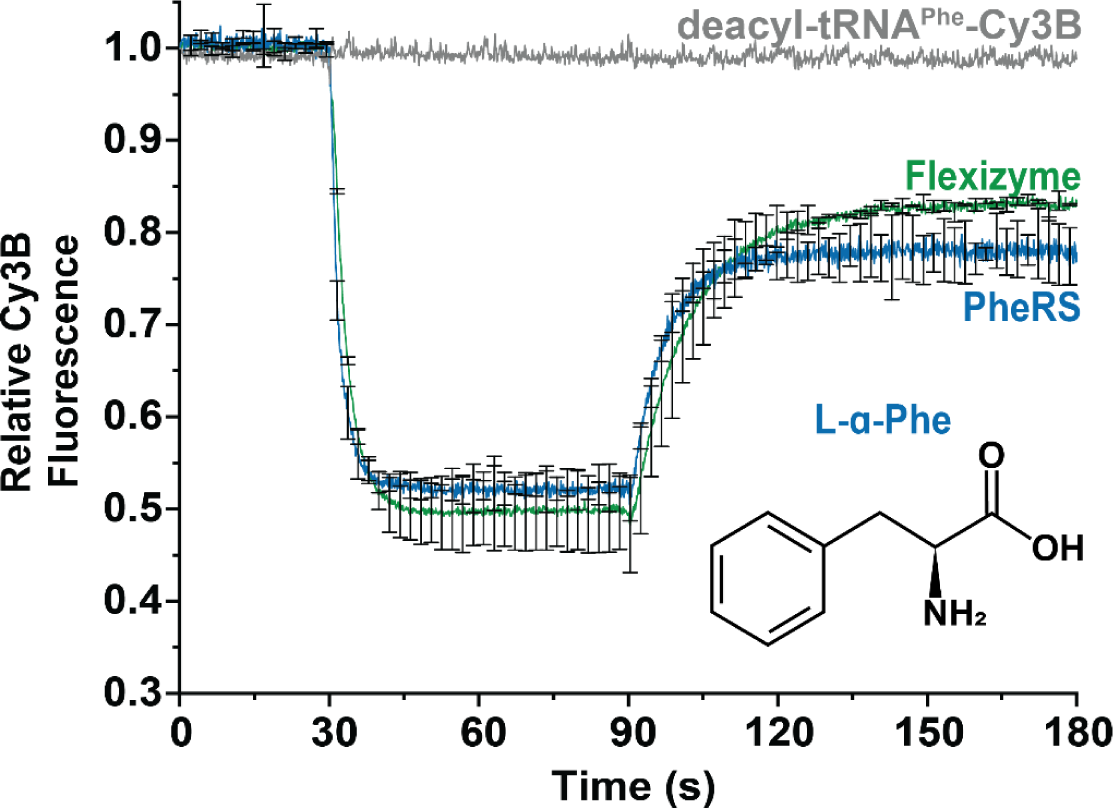
Ternary complex formation assay comparing Flexizyme and PheRS-charged L-α-Phe-tRNA^Phe^-Cy3B. Ternary complex formation assay as described in Figure 2 comparing tRNA^Phe^-Cy3B charged by PheRS (blue) or flexizyme (green), and de-acylated tRNA^Phe^-Cy3B (grey). Error bars represent S.D. from two separate replicates.

**Supplemental Figure S3.**
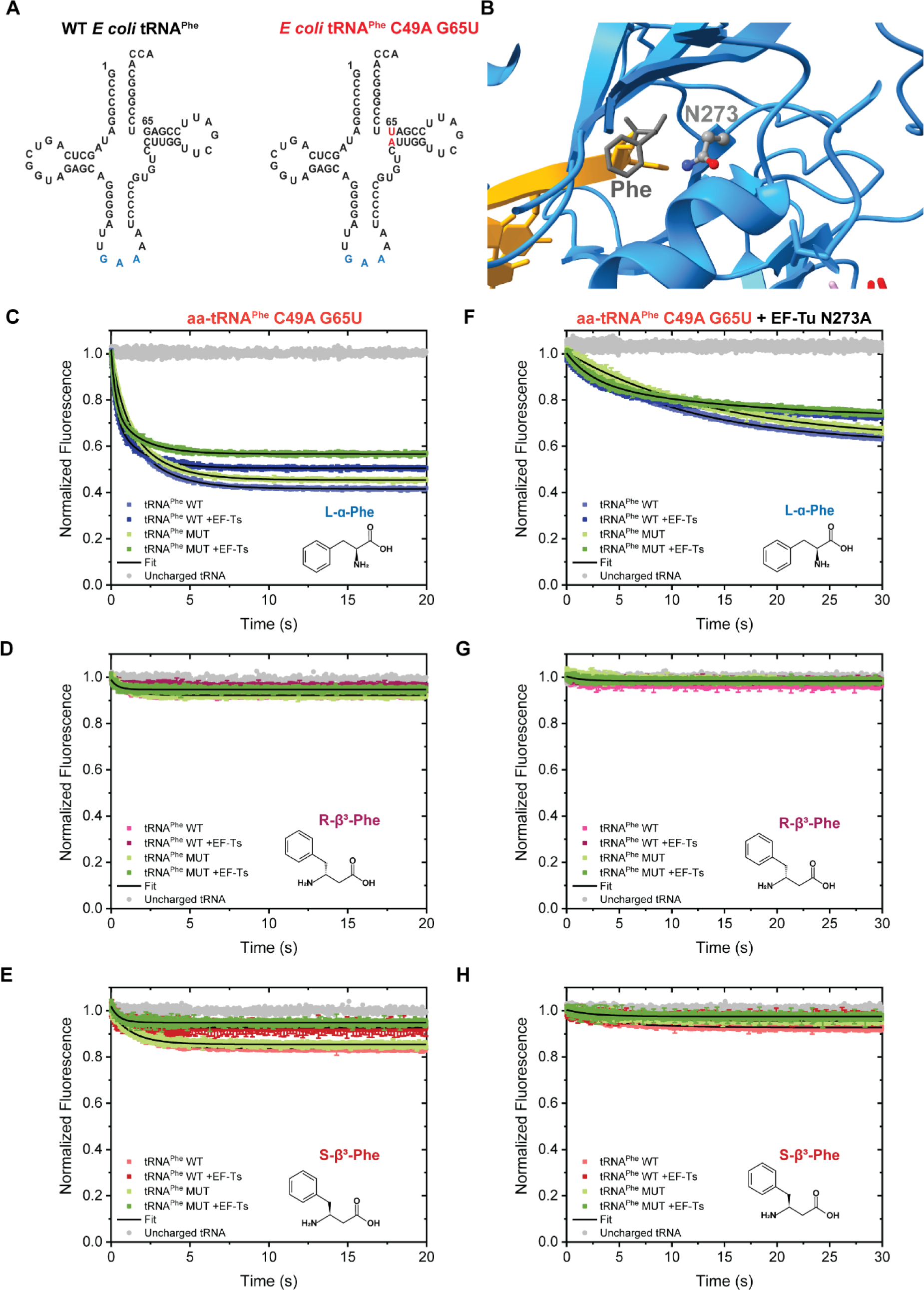
Mutant tRNA^Phe^ (C49A G65U) and/or EF-Tu (N273A) examined by stopped-flow ternary complex formation. **A)** Sequences of WT (left) and mutant (right) *E coli* tRNA^Phe^ used for stopped-flow kinetic analysis. **B)** Structure of EF-Tu (PDB: 1OB2) with the N273 amino acid residue in the amino acid binding pocket shown next to the Phe aminoacylated to the A76 on the 3’-end of tRNA^Phe^ (yellow). **C-E)** Stopped-flow ternary complex formation assays using Cy3B-labeled mutant aa-tRNA^Phe^ (MUT) with the indicated monomers inset. **F-H)** Stopped-flow ternary complex assays with both tRNA^Phe^ MUT and mutant EF-Tu N273A with the indicated monomers inset. Error bars represent S.D. of 3-5 replicates.

**Supplemental Figure S4.**
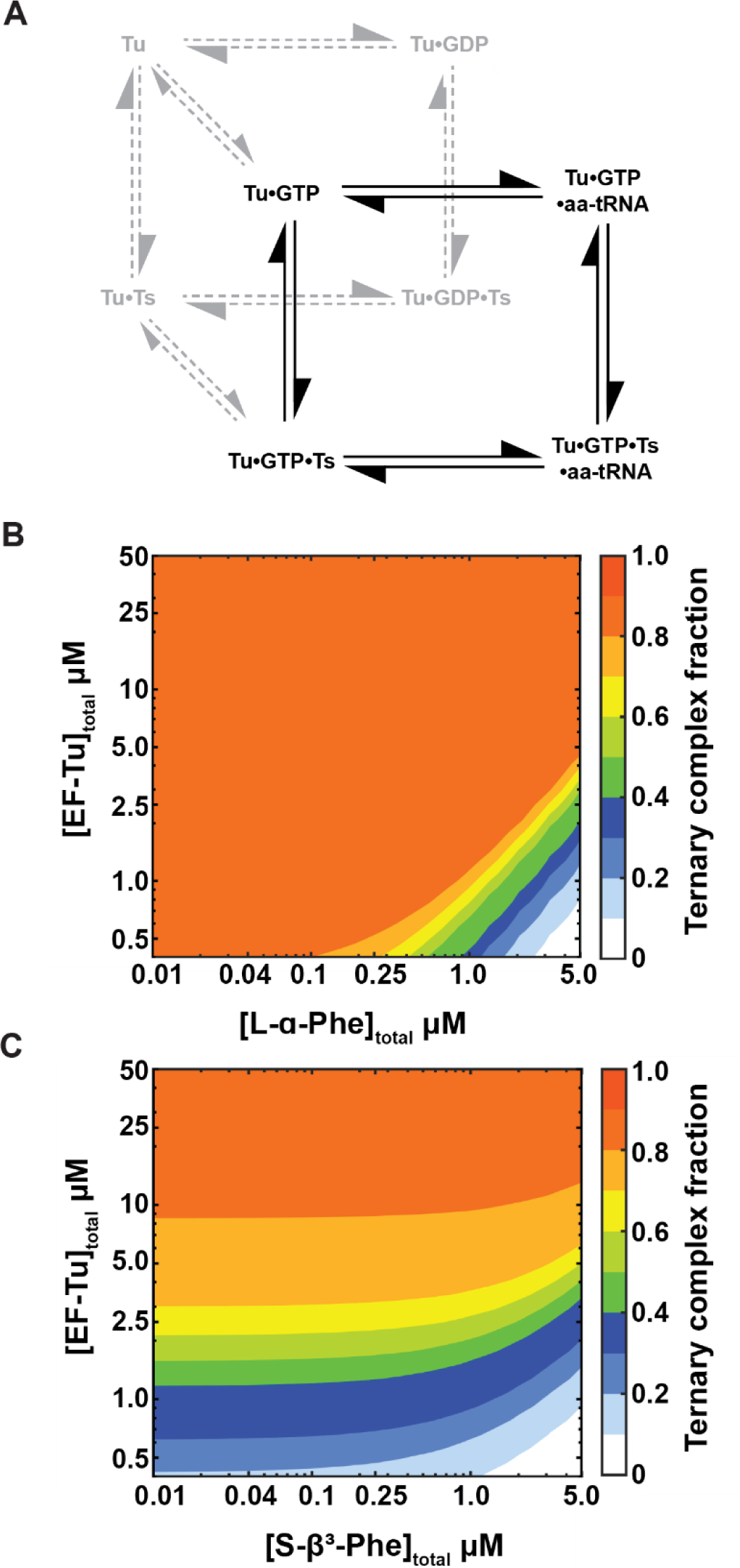
Kinetic simulations of ternary complex formation with S-β^3^-Phe. **A)** Minimal reaction scheme for kinetic simulations. Contour plots of **B)** L-α-Phe and **C)** S-β^3^-Phe ternary complex abundance simulated at different physiologically relevant aa-tRNA and ET-Tu/Ts concentrations. Simulations were done using measured rate constants from experiments reported here and from the literature. Ternary complex fraction was calculated as [aa-tRNA]_bound_/[aa-tRNA]_total_

**Supplemental Figure S5.**
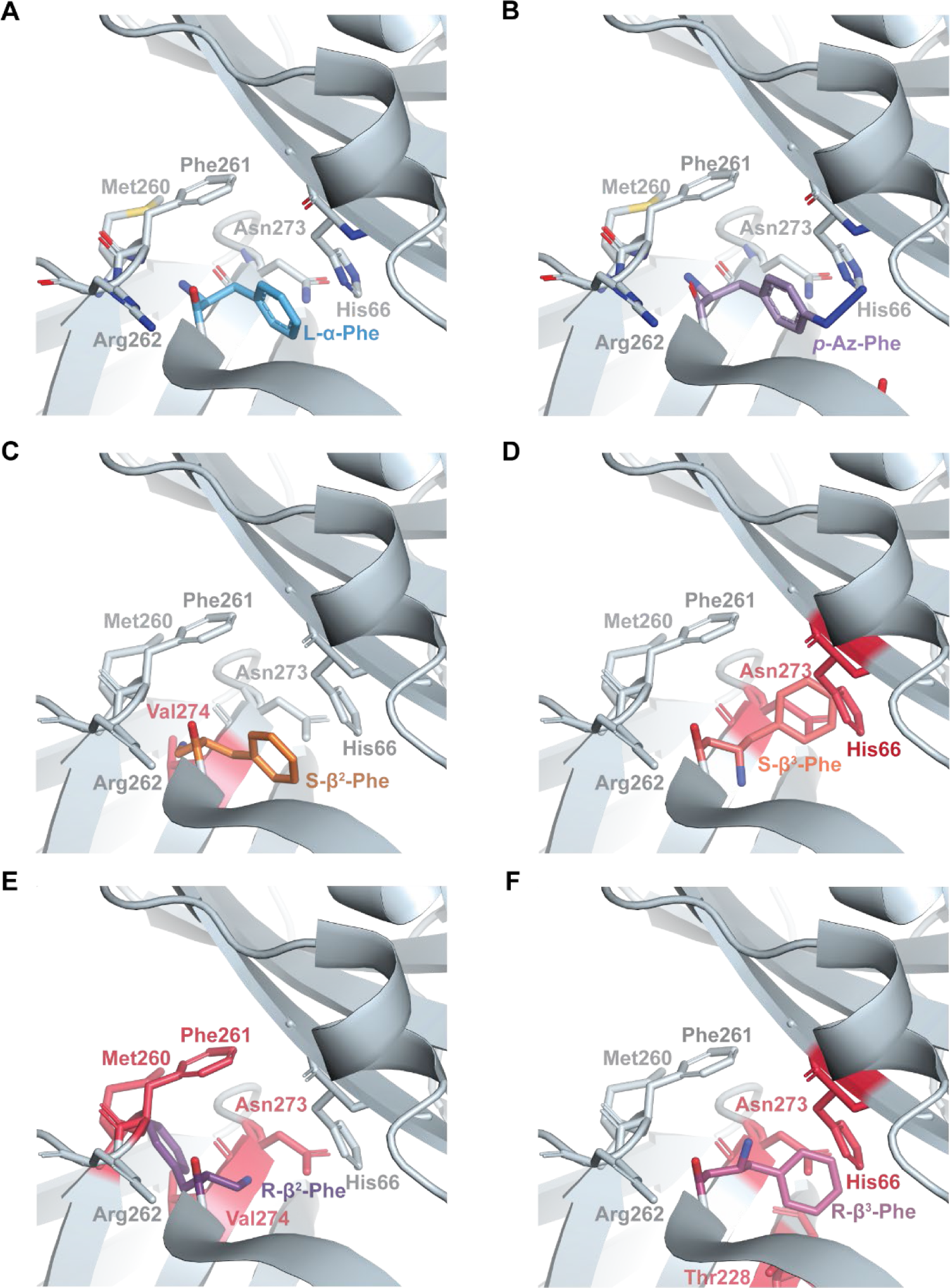
Manual modelling of aa-tRNA^Phe^ into the EF-Tu binding pocket. Manual modelling of **A)** L-α-Phe, **B)** *p*-Az-Phe, **C)** S-β^2^-Phe, **D)** S-β^3^-Phe, **E)** R-β^2^-Phe, **F)** R-β^3^-Phe into the amino acid binding pocket of EF-Tu formed by the DI-DII interface. Steric clashes are represented in red. The 2D model of non-natural amino acids was created using the Chemical Sketch Tool provided by the RCSB Protein Data Bank. Subsequently, the 3D atomic models and RNA-peptide links were constructed using JLigand ^80^. The figures were generated using PYMOL, the PyMOL Molecular Graphics System, Version 2.5.7 Schrödinger, LLC.

---

**TOC Figure**

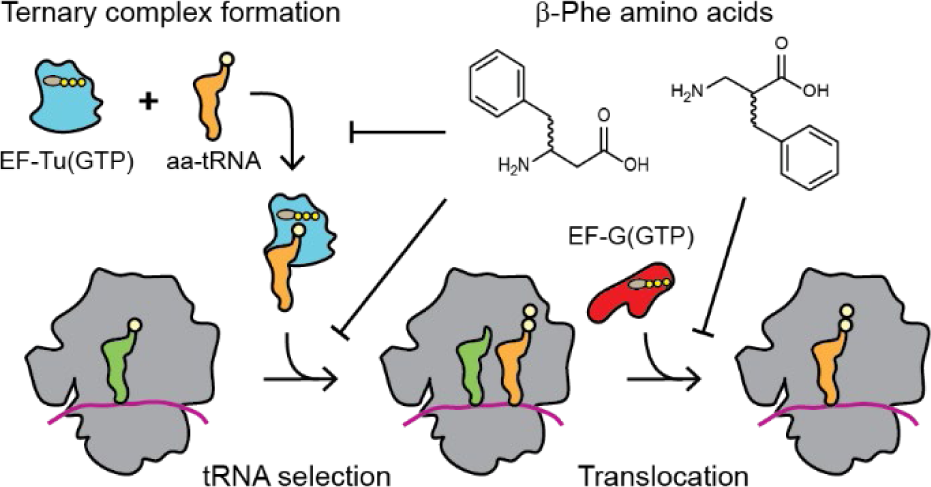

